# HIV-SEQ Reveals Global Host Gene Expression Differences Between HIV-Transcribing Cells from Viremic and Suppressed People with HIV

**DOI:** 10.1101/2024.12.17.629023

**Authors:** Julie Frouard, Sushama Telwatte, Xiaoyu Luo, Natalie Elphick, Reuben Thomas, Douglas Arneson, Pavitra Roychoudhury, Atul J Butte, Joseph K Wong, Rebecca Hoh, Steven G. Deeks, Sulggi A. Lee, Nadia R. Roan, Steven Yukl

**Affiliations:** Gladstone Institutes, San Francisco, CA 94158, USA; Department of Urology, University of California, San Francisco, CA 94158, USA; San Francisco Veterans Affairs (VA) Medical Center and University of California, San Francisco, CA, USA; Department of Infectious Diseases, The University of Melbourne at the Peter Doherty Institute of Infection and Immunity, Melbourne, Australia; Bakar Computational Health Sciences Institute, University of California, San Francisco, San Francisco, CA, USA; Department of Laboratory Medicine and Pathology, University of Washington, Seattle, Washington, USA; Viral and Infectious Disease Division, Fred Hutchinson Cancer Research Center, Seattle, Washington, USA; Division of HIV, Infectious Diseases and Global Medicine, University of California, San Francisco, USA; Zuckerberg San Francisco General Hospital and the University of California, San Francisco, USA

## Abstract

“Active” reservoir cells transcribing HIV can perpetuate chronic inflammation in virally suppressed people with HIV (PWH) and likely contribute to viral rebound after antiretroviral therapy (ART) interruption, so they represent an important target for new therapies. These cells, however, are difficult to study using single-cell RNA-seq (scRNA-seq) due to their low frequency and low levels of HIV transcripts, which are usually not polyadenylated. Here, we developed “HIV-seq” to enable more efficient capture of HIV transcripts – including non-polyadenylated ones – for scRNA-seq analysis of cells from PWH. By spiking in a set of custom-designed capture sequences targeting conserved regions of the HIV genome during scRNA-seq, we increased our ability to find HIV RNA+ cells from PWH by up to 44%. Implementing HIV-seq in conjunction with surface phenotyping by CITE-seq on paired blood specimens from PWH before vs. after ART suppression, we found that HIV RNA+ cells were enriched among T effector memory (Tem) cells during both viremia and ART suppression, but exhibited a cytotoxic signature during viremia only. By contrast, HIV RNA+ cells from the ART-suppressed timepoints exhibited a distinct anti-inflammatory signature involving elevated TGF-β and diminished IFN signaling. Overall, these findings demonstrate that active reservoir cells exhibit transcriptional features distinct from HIV RNA+ cells during viremia, and underscore HIV-seq as a useful tool to better understand the mechanisms by which HIV-transcribing cells can persist during ART.

## INTRODUCTION

Most people with HIV (PWH) experience rebound of HIV in plasma within several weeks after stopping antiretroviral therapy (ART), indicating the persistence of a “rebound-competent” viral reservoir that prevents HIV cure. The prevailing model has been that the rebound virus arises from a small fraction of HIV-infected cells that contain an infectious provirus in a latent state, where the infected cell does not constitutively produce virions but can be induced to do so after activation^1–3^. However, the rebound virus is often different from that which grows out *ex vivo* following stimulation in quantitative viral outgrowth assays (QVOA)^4,5^, suggesting that additional studies are needed to understand the reservoirs which can be reactivated *in vivo*. A smaller body of research has focused on the cells actively transcribing HIV RNA *in vivo*, also known as the “active” reservoir. While HIV latency and expression are often viewed as a dichotomy (off/on), studies using multiple round QVOAs demonstrate varying degrees of inducibility *ex vivo*^6^, and blood and tissue cells from ART-suppressed individuals in fact show a continuum of HIV expressing cells *in vivo*, with variable degrees of progression through different blocks to HIV expression^7–9^.

Importantly, multiple lines of evidence suggest that HIV-infected cells which spontaneously express HIV RNA or protein *in vivo* may be just as important for pathogenesis and cure as the transcriptionally-silent reservoir. First, viral products expressed by active reservoir cells are likely to contribute to the immune activation and inflammation^10,11^ that are thought to underlie the sequelae of ART-treated infection, including organ damage and reduced life expectancy^12–16^. Second, the active reservoir seems better poised than the silent reservoir to immediately infect new cells after interruption of ART because it does not need to revert the mechanisms (e.g. epigenetic modifications) that prevent viral expression in the silent reservoir. Indeed, at least four studies have shown that different forms of cell-associated HIV RNA negatively correlate with time to rebound after ART treatment interruption (ATI)^17–20^. Moreover, one small study showed that in about half of people who interrupted ART, Pol sequences from the rebound virus matched those from cell-associated HIV RNA prior to ART interruption^21^. These findings support the hypothesis that the active reservoir is rebound-competent. Finally, active reservoir cells, by expressing HIV RNA and/or protein, are likely more susceptible than quiescent latent cells to new immune-based therapies aimed at an HIV cure (e.g. therapeutic vaccines, TLR agonists, adoptive immune therapies, and broadly neutralizing antibodies).

Despite all this evidence pointing towards the active reservoir as an important target for HIV cure, our understanding of these cells is still rudimentary, and it is unclear whether these cells exhibit similar phenotypic features as HIV RNA+ cells present during viremia. Cells transcribing or translating HIV genes from viremic individuals have been characterized by flow cytometry, CyTOF, and single-cell sequencing^22–25^ to a certain extent, but HIV-transcribing cells from ART-suppressed PWH have been harder to study with such single-cell technologies. For instance, conventional sequencing-based approaches identify active reservoir cells at such low numbers as to preclude meaningful analysis^25^. One alternative has been to activate cells from ART-suppressed individuals *ex vivo* to characterize reactivated HIV RNA+ cells^26,27^, but this approach characterizes *ex vivo* stimulated and not spontaneously HIV-transcribing cells. Of note, sequencing-based approaches for detection of HIV RNA+ cells from virally-suppressed PWH are not only limited by the low throughput and high costs of droplet-based encapsulation technologies (e.g. 10X Genomics), but also the reliance on RNA capture through poly-dT. Because most HIV-infected cells from ART-suppressed individuals do not contain polyadenylated HIV RNA due to blocks to transcriptional elongation and completion^7^, poly-dT-based methods of RNA capture will theoretically fail to recognize many HIV-transcribing cells.

To increase the ability to identify rare, HIV RNA+ cells from PWH, including in the context of ART suppression, we created a custom-modified, 10x Genomics-compatible, 5’ sequencing-based scRNA-seq workflow in which the poly-dT primer is supplemented with multiple HIV-specific reverse primers targeting different regions of the genome. This approach captures HIV-infected cells with non-polyadenylated HIV transcripts, including 5’ elongated as well as more distal HIV transcripts. In addition, we included DNA-barcoded antibodies (CITE- seq^28^) in our protocol to enable in-depth phenotyping of the HIV RNA+ cells for which we have transcriptome data. Compared to the standard 5’ sequencing approach, the inclusion of HIV-specific reverse primers allowed the detection of more HIV RNA+ cells from PWH, and enabled for the first time a meaningful analysis of HIV RNA+ cells from ART-suppressed PWH. Using this advantage, we performed an in-depth analysis of the transcriptomes and phenotypes of HIV RNA+ cells from longitudinal samples of PWH during active viremia and after ART suppression.

## RESULTS

### Development of HIV-seq as a method to increase capture and detection of HIV transcripts by single-cell sequencing

The standard 10X Genomics’ 5’ scRNA-seq workflow entails droplet encapsulation of individual cells, followed by capture and reverse transcription of polyadenylated transcripts using poly(dT) oligos. This approach, theoretically, does not capture non-polyadenylated transcripts. To efficiently capture HIV transcripts, including those that are not poly-adenylated, we designed capture sequences targeting multiple conserved regions of the HIV genome (Fig. 1A): the R-U5-pre-Gag region (for 5’ elongated transcripts), the Pol gene (for mid-transcribed, unspliced transcripts), the second exon of Tat-Rev (for distal transcripts, including spliced), and two regions known to be enriched among intact proviruses: the packaging signal^29^ (Psi; for elongated, unspliced transcripts) and the HIV Rev response element^29^ (RRE, for distal and unspliced/single-spliced transcripts). Our capture sequences match the consensus sequence of subtype B HIV-1, and are known to be efficient reverse primers for established ddPCR assays^7^. The HIV capture oligos were spiked into the poly-dT primer mix and used in the reverse transcription (RT) following cell encapsulation (Fig. 1B). After RT, gene expression and CITE-seq libraries were prepared and sequenced. We aligned all sequences to the GRCh38 human genome, to which we had appended a subtype B consensus sequence we had generated from the Los Alamos database (see Methods). We named our overall pipeline “HIV-seq” due to its specific targeting of HIV transcripts for RT, library generation, and alignment.

**Figure 1.**
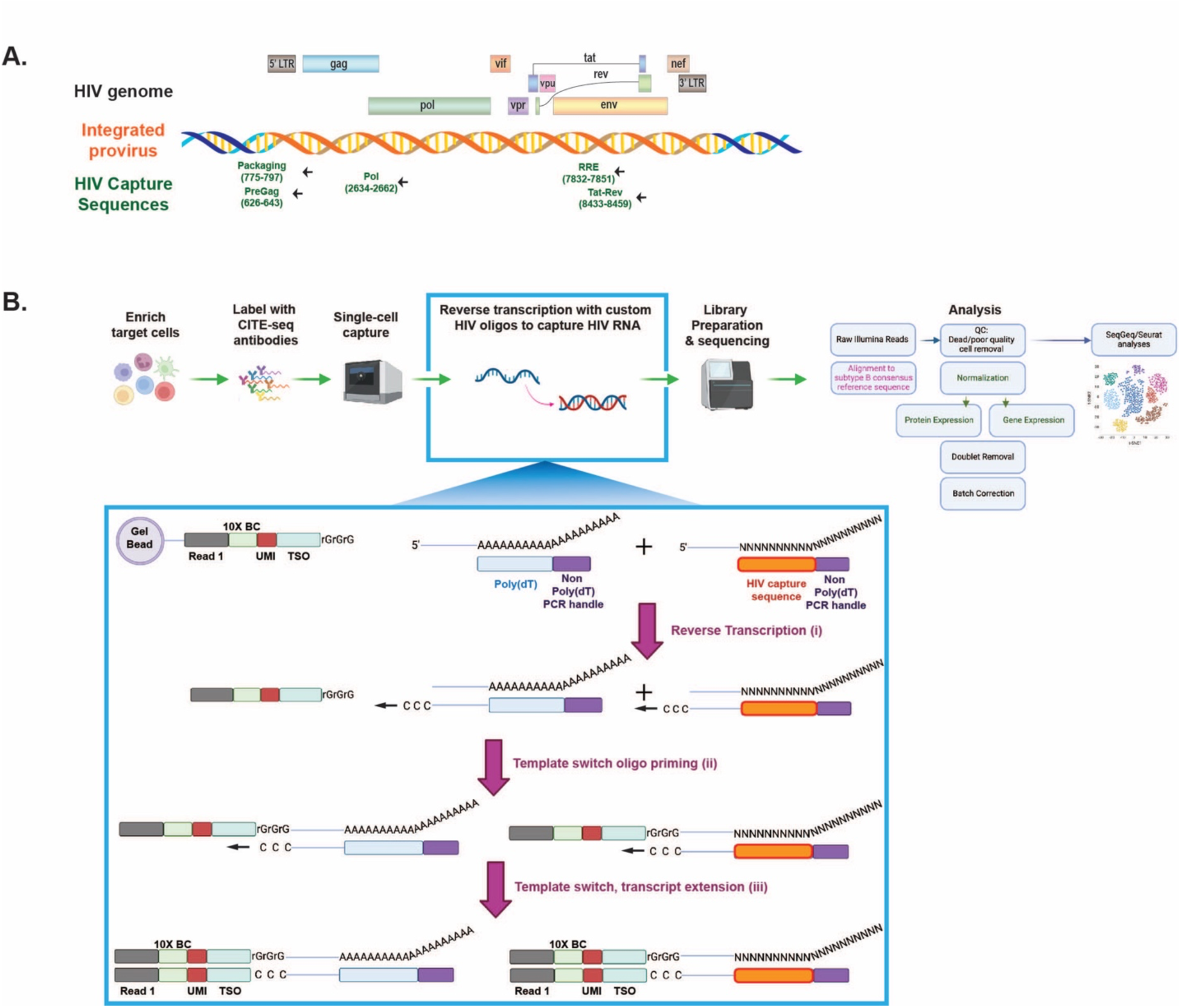
HIV-seq method to increase detection of HIV transcripts by single-cell sequencing. **A.** Schematic illustrating the genomic location of HIV-specific capture sequences of HIV-seq. The name and nucleotide position (based off HXB2 annotation) of each of capture sequence are labeled in green. **B.** Schematic of HIV-seq protocol. PBMCs from PWH are enriched for CD4+ T cells and then labeled with CITE-seq antibodies to enable subsequent surface phenotyping. After cell encapsulation using a 10X Chromium instrument, custom-designed HIV-specific capture sequences described in *panel A* that have been appended to a non-poly(dT) PCR handle are spiked in with the poly(dT) oligos and incorporated into the 10X Genomics’ Chromium Next GEM Single Cell 5’ workflow. Each ‘Single Cell 5’ Gel Bead’ features an Illumina R1 sequence (‘read 1’ sequencing primer), a 16 nucleotide (nt) 10X Barcode (BC), a 10 nt unique molecular identifier (UMI), and a 13 nt template switch oligo (TSO). Reverse transcription is primed off both poly(dT) as well as the HIV capture sequences (i). Following template switching and transcript extension (ii, iii), barcoded cDNA libraries corresponding to both gene expression (GEX) and antibody-derived tags (ADT, for CITE-seq) are processed through the standard 10X workflow, and then sequenced. Data analysis was performed using the Seurat pipeline, SeqGeq software, and custom scripts.

To assess the utility of HIV-seq, we compared it head-to-head to the original 10X Genomics’ 5’ single cell RNA-seq pipeline (without HIV primers). To obtain sufficient numbers of HIV RNA+ cells for this comparative analysis, we selected samples from two viremic donors (PID1052 and PID8027) not on ART. Similar numbers of cells were processed using the conventional vs. the HIV-seq pipeline and then analyzed by scRNA-seq. We first confirmed that in both donors, there was no difference in global gene expression between the two experimental conditions (Fig. 2A), thereby demonstrating that HIV-seq does not perturb the capture and sequencing of the host transcriptome. It also does not result in spurious detection of HIV RNA transcripts in non-permissive CD8+ T cells (Fig. S1). Quantitative analyses showed that HIV-seq identified a higher percentage of HIV RNA+ cells from both donors than did traditional 5’ scRNA-seq (Fig. 2B). For PID1052, HIV-seq increased the identification of HIV RNA+ cells from 0.047% to 0.068% of the CD4+ T cell population, corresponding to a 44.7% increase. For PID8027, who had more HIV RNA+ cells, HIV-seq increased the capture of HIV RNA+ cells from 0.82% to 0.97%, corresponding to an 18.3% increase. The numbers of HIV transcripts detected per infected cell also increased significantly with HIV-seq, from a mean of 22 to 44 reads/cell (Fig. 2B, 2C).

**Figure 2.**
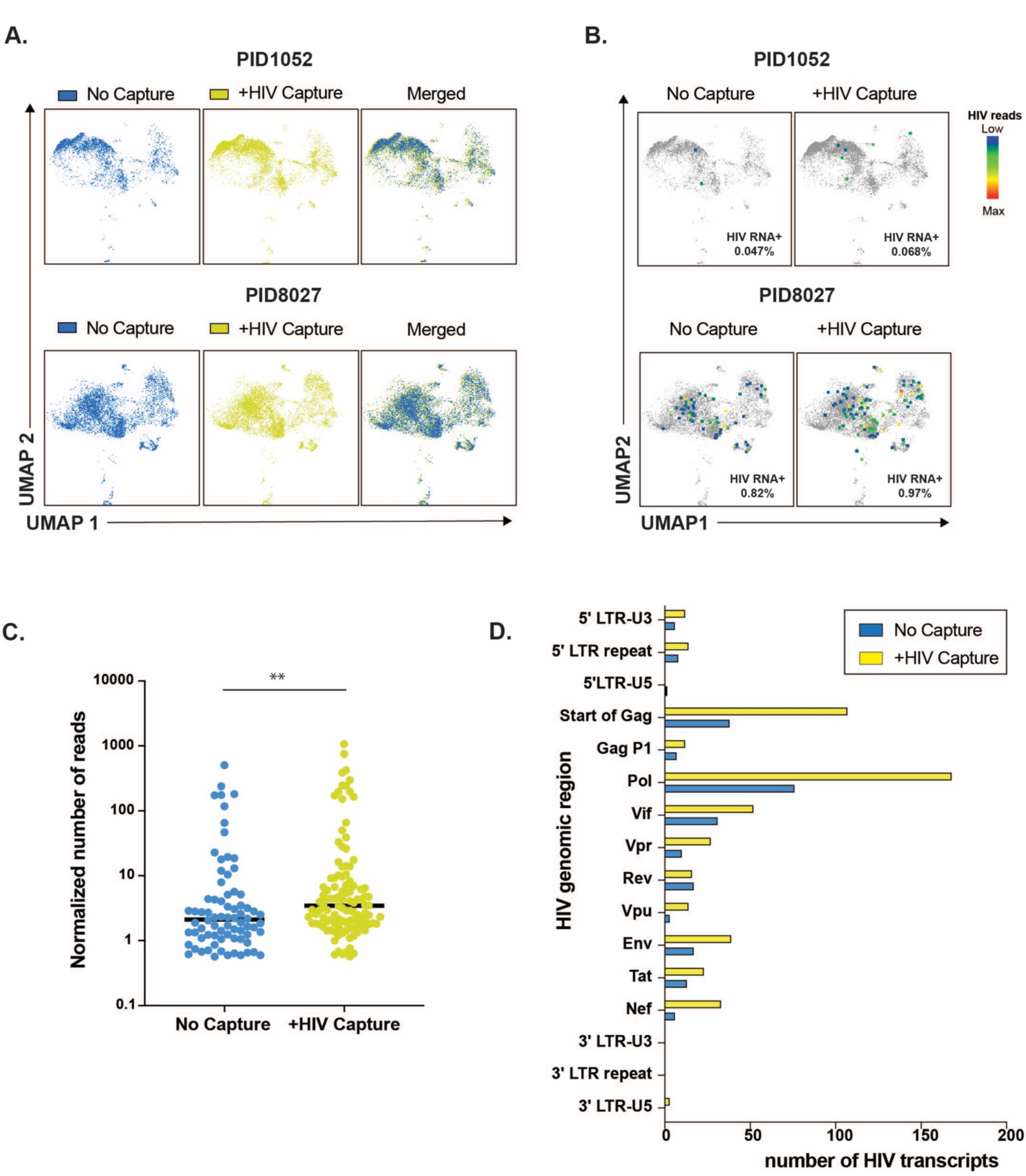
HIV-seq increases detection of HIV RNA+ cells from PWH without affecting host transcriptome. **A.** HIV-seq does not affect host transcriptome analysis by scRNAseq. Shown are UMAP plots of CD4+ T cells from two ART-naïve viremic donors (PID1052 and PID8027), processed in the absence (blue) vs. presence (yellow) of HIV capture sequences. **B.** HIV-seq increases numbers of HIV RNA+ cells identified from viremic PWH. UMAP plots of scRNAseq analysis of pre-ART CD4+ T cells from PID1052 and PID8027, showing HIV RNA+ cells as colored dots among total CD4+ T cells represented in gray, with vs. without the addition of HIV capture sequences. The percentages of HIV RNA+ cells among CD4+ T cells are indicated in the lower right of each plot. Colors represent the number of HIV reads, from low (blue) to high (red). **C.** HIV-seq increases the numbers of HIV RNA reads detected per infected cell. Normalized numbers of HIV reads in the absence vs. presence of HIV capture sequences, for each HIV RNA+ cell identified from PID1052 and PID8027 in *panel A*. Horizontal lines correspond to median values. **P ≤ 0.01 as determined using a Mann-Whitney test. **D.** HIV-seq increases detection of HIV reads throughout the HIV genome. HIV reads from PID1052 and PID8027 were aligned to the HIV-1 subtype B consensus reference genome. The y-axis depicts individual HIV genes, and the x-axis shows the number of detected HIV transcripts, in the absence (blue) vs. presence (yellow) of the HIV capture sequences.

To assess whether the additional HIV transcripts captured by HIV-seq preferentially mapped to certain regions of the HIV genome, we compared the distribution of the HIV reads from the two viremic individuals in the absence vs. presence of HIV-capture oligos. HIV-seq increased the detection of HIV transcripts from across the entire proviral genome, without changing the representation of the HIV regions detected relative to traditional 5’ scRNA-seq (Fig. 2D). Under both conditions, most HIV transcripts aligned to the *pol* and early *gag* regions of HIV. Application of HIV-seq in the context of ART suppression also resulted in detection of transcripts primarily mapping to the *gag* and *pol* regions (Fig. S2). Overall, these results demonstrate that HIV-seq enables more efficient detection of HIV RNA+ cells and of HIV transcripts per infected cell than does conventional 5’ scRNA-seq.

### HIV RNA+ cells from viremic PWH are preferentially cytolytic Tem cells and exhibit an intracellular state promoting HIV replication

Leveraging the ability of HIV-seq to increase the numbers of HIV RNA+ cells we can analyze by scRNA-seq, we implemented it on cells from 4 viremic, ART-naïve PWH (PID1052, PID8026, PID8027, and PID0145). In total, 1,072 HIV RNA+ CD4+ T cells were identified, and infected cell frequencies ranged from 0.061% to 2.42% of the CD4+ T cell population (Fig. 3A). UMAP visualization of the transcriptomes of the HIV RNA+ cells revealed heterogeneity, in that infected cells were distributed in multiple regions of the UMAP space, yet enrichment was observed in some regions, suggesting non-random distribution of infected cells among CD4+ T cell subsets (Fig. 3B). Considerable variability in the degree of HIV transcription was observed among infected cells, with HIV transcript levels ranging from 1 to 1063 HIV reads per cell. When we separated the HIV RNA+ cells into those with low numbers of HIV reads (HIV_low_: 1 to 50 HIV reads) and those with high numbers of HIV reads (HIV_high_: > 50 HIV reads), we found that participants with lower numbers of HIV RNA+ cells (PID1052 and PID0145) only harbored HIV_low_ cells, while those with higher numbers (PID8026 and PID8027) had both populations (Fig. S3A). Of note, HIV-seq did not affect whether HIV_high_ could be detected (Fig. S3B). The distribution of HIV_high_ and HIV_low_ cells across the UMAP was similar (Fig. S3C) and the only significantly differentially expressed transcript/protein between these populations – apart from HIV RNAs – was the CD4 protein, which was decreased among the HIV_high_ cells (Fig. S3D).

**Figure 3.**
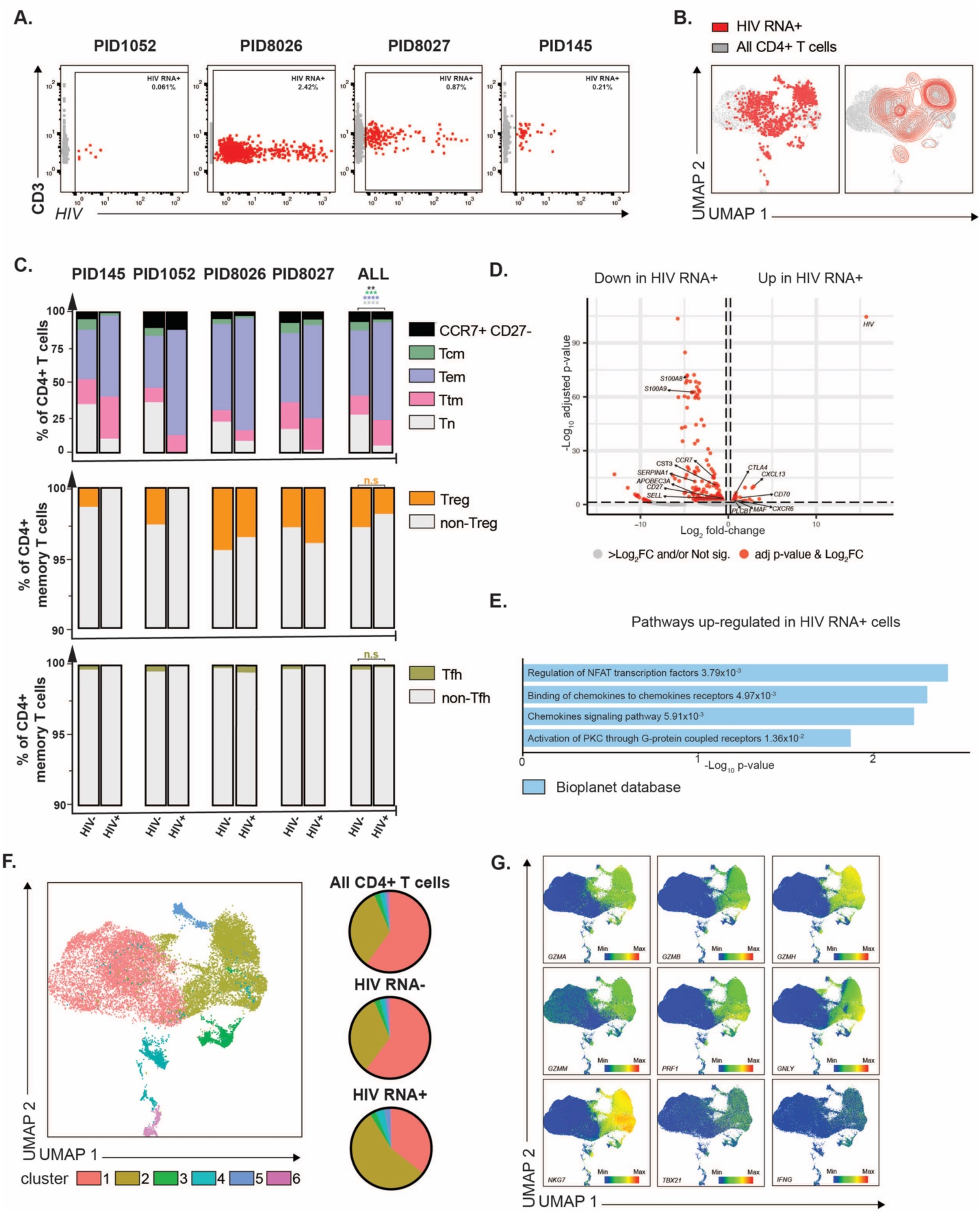
HIV RNA+ cells from viremic PWH are preferentially Tem cells and exhibit transcriptional signatures of cytolysis and cellular activation. **A.** CD3+CD4+ T cells transcribing HIV RNA were identified by HIV-seq from 4 viremic PWH (PID1052, PID8026, PID8027 and PID145) based on HIV RNA levels. HIV RNA+ cells are depicted in red and HIV RNA- cells are depicted in gray. Percentages of HIV RNA+ cells are indicated in the upper right of each plot. **B.** A diverse array of HIV RNA+ cells exists among CD4+ T cells from viremic PWH. Shown are UMAP plots depicting HIV RNA+ cells as red dots (*left*) or contours (*right*), against a background of HIV RNA- CD4+ T cells depicted in gray, for all 4 donors listed in *panel A* combined. **C.** HIV RNA+ cells from viremic PWH are enriched in T effector memory (Tem) cells and disenriched in T central memory (Tcm) and naïve T (Tn) cells relative to HIV RNA- CD4+ T cells (top panel). The proportion of Treg vs non-Treg (middle panel) and Tfh vs non-Tfh cells (bottom panel) is not significantly different in HIV RNA+ vs HIV RNA- memory CD4+ T cells. **P ≤ 0.01, ***P≤ 0.001, ****P≤ 0.0001 as determined by generalized linear mixed model (GLMM). **D.** HIV RNA+ cells differentially express host transcripts involved in immune responses. Shown is a volcano plot displaying upregulated and downregulated transcripts in HIV RNA+ vs. HIV RNA- CD4+ T cells, with select transcripts annotated. Red dots correspond to transcripts with 0.25log_2_ fold-change expression and with an adjusted p value ≤ 0.05, as determined by genewise quasi F-tests. **E.** HIV RNA+ cells exhibit gene expression signatures of cellular activation, inflammation, and chemokine signaling. Shown is a pathway analysis comparing HIV RNA+ HIV RNA- cells, using the Bioplanet 2019 database. The p-value representing the statistical significance of enrichment of the gene set within the pathway is indicated on the right. **F.** HIV RNA+ cells are enriched in cluster 2 of CD4+ T cells. *Left:* UMAP depicting the 6 clusters of CD4+ T cells identified by Louvain clustering. *Right:* Pie graphs showing the distribution of total CD4+ T cells (*top*), HIV RNA- CD4+ T cells (*middle*) or HIV RNA+ CD4+ T cells (*bottom*) among the different clusters. **G.** Cluster 2 cells, concentrated on the right of the UMAP, are defined by high expression of cytotoxic markers *GZMA, GZMB, GZMH, GZMM, PRF1, GNLY, NKG7*, and Th1-defining factors *IFNG* and *TBX21*. Heatmaps depict relative expression of the indicated transcripts.

This finding likely reflects higher expression of Nef, which downregulates cell-surface CD4^30^, in the HIV_high_ cells. Because the HIV_high_ and HIV_low_ cells exhibited overall similar gene expression profiles, all remaining analyses combined these two populations together.

We then assessed whether the HIV RNA+ cells were enriched in specific cellular subsets. We first assessed the distribution of classical CD4+ T cell subsets (Fig. S4) among uninfected and HIV RNA+ CD4+ T cells. Relative to their uninfected CD4+ T cell counterparts, HIV RNA+ cells were under-represented among naïve cells and over-represented among memory cells (Fig. 3C), as expected^31^. Within memory T cells, HIV RNA+ cells were under- represented among central memory (Tcm) cells and those of the CCR7+CD27- phenotype, and were over-represented among effector memory (Tem) cells (Fig. 3C), consistent with prior reports of over-representation of Tem among HIV RNA+ cells from viremic individuals^25,32^. By contrast, transitional memory (Ttm), regulatory T cells (Treg), and T follicular helper (Tfh) cells were equally represented among uninfected and HIV RNA+ CD4+ T cells (Fig. 3C).

Next, we compared differentially expressed transcripts between HIV RNA+ and HIV RNA- cells from the viremic PWH. Almost 300 genes were differentially expressed (Fig. 3D, Table S3). Consistent with disenrichment of Tcm among HIV RNA+ cells (Fig. 3C), transcript levels of the *CCR7*, *SELL* (encoding the protein CD62L), and *CD27* – markers of Tcm cells^33,34^ – were downregulated in the HIV RNA+ cells (Fig. 3D). HIV RNA+ cells also had low transcript levels of the alarmins *S100A8* and *S100A9*, which encode for S100 calcium binding protein A8 and A9, respectively. These proteins are released in response to environmental triggers and cellular damage^35^. Antiviral factors, including *SERPINA1*^36,37^ and *APOBEC3A*^38^, were also decreased among HIV RNA+ cells, suggesting that the intracellular state of HIV RNA+ cells in viremic individuals may favor HIV replication. In line with this finding, HIV RNA+ cells also exhibited downregulation of *CST3*, which encodes cystatin C, a cysteine protease inhibitor that interacts with the HIV proteins gp160, gp120, p31 and p24, and inhibits HIV protease function^39^. With regards to transcripts upregulated among HIV RNA+ cells, HIV transcripts were the top hit, as expected. In addition, HIV RNA+ cells expressed higher levels of the activation marker *CD70*. Interestingly, CD4+CD70+ cells are increased in PWH with high levels of viremia and associate with immune activation^40^, suggesting that this subset of infected cells may contribute to disease progression during untreated infection. HIV RNA+ cells also expressed elevated levels of *CXCR6*, which encodes a chemokine receptor that is an alternative co-receptor for HIV^41^.

Pathway analysis of the DEGs revealed elevated upregulation of the NFAT pathway (*MAF*, *CLTA4*) in HIV RNA+ cells (Fig. 3D, 3E), consistent with NFAT as a driver of HIV transcription^42,43^. Similarly, there was an upregulation of the PKC pathway, known for its involvement in HIV gene expression and latency reversal^44^. Finally, and consistent with the DEG analysis, chemokine signaling pathways – featuring genes such as *CXCR6*, the CXCR6 ligand *CXCL13*, and *PLCB1* – were also elevated among HIV RNA+ cells.

While DEG analysis identified both known and novel shared features among all HIV RNA+ cells, it was clear that the HIV RNA+ cells were heterogeneous (Fig. 3B). We therefore implemented clustering to assess for transcriptomic or phenotypic features that may not be shared by the entire population of infected cells. Louvain clustering identified six clusters of CD4+ T cells (Fig. 3F). The classical CD4+ T cell subsets (Fig. S4) did not define the clusters, and in fact all the classical memory subsets were represented among the 6 clusters, albeit in different proportions (Fig. S5). The HIV RNA+ cells were differentially distributed among the six clusters as compared to their uninfected counterparts, with enrichment of HIV RNA+ cells among cluster 2 (Fig. 3F). This cluster was characterized by high expression of cytotoxic and cytolytic genes, including *GZMA, GZMB, GZMH, GZMM, PRF1, GNLY, NKG7,* as well as the Th1-defining factors *IFNG* and *TBX21* (Fig. 3G). This finding suggests that cluster 2 cells are of a cytolytic Th1 phenotype, and is consistent with prior reports that HIV RNA+ cells from viremic PWH exhibit Th1 cytolytic signatures^25,32^.

Overall, our data demonstrate that HIV RNA+ cells from viremic PWH are heterogeneous but exhibit shared features, including being more likely to be Tem, and notably, displaying a previously undescribed state more conducive to viral replication, with low expression of restriction factors and increased activation of cellular pathways promoting HIV gene expression. Additionally, relative to their uninfected counterparts, they preferentially exhibit a cytolytic signature characterized by higher expression of granzymes, perforin, and granulysin and a Th1 signature.

### HIV RNA+ cells from suppressed PWH are preferentially Tem cells but do not exhibit a cytolytic signature

Three of the individuals we had analyzed in the context of active viremia (PID1052, PID8026, and PID8027) had specimens collected after > 24 weeks of ART suppression.

Therefore, we next implemented a similar HIV-seq analysis pipeline to characterize these participant-matched HIV RNA+ cells in the context of ART suppression. A total of 26 HIV RNA+ cells were detected, corresponding to an infected cell frequency of 0.016% to 0.091% (Fig. 4A). As for the viremic specimens, HIV RNA+ cells were broadly distributed among CD4+ T cells, demonstrating heterogeneity (Fig. 4B), and enriched among Tem cells (Fig. 4C); in fact, in two individuals (PID8026 and PID8027), the entire HIV RNA+ cell population was exclusively found within this subset. Few DEGs were observed between the HIV RNA+ and HIV RNA- CD4+ T cells, with the notable exception of a handful of downregulated genes (Fig. 4D and Table S4). These included the Tcm marker *SELL,* the alarmin *S100A9*, and the cysteine protease inhibitor *CST3*, all of which were also preferentially downregulated among HIV RNA+ cells during active viremia (Fig. 3).

**Figure 4.**
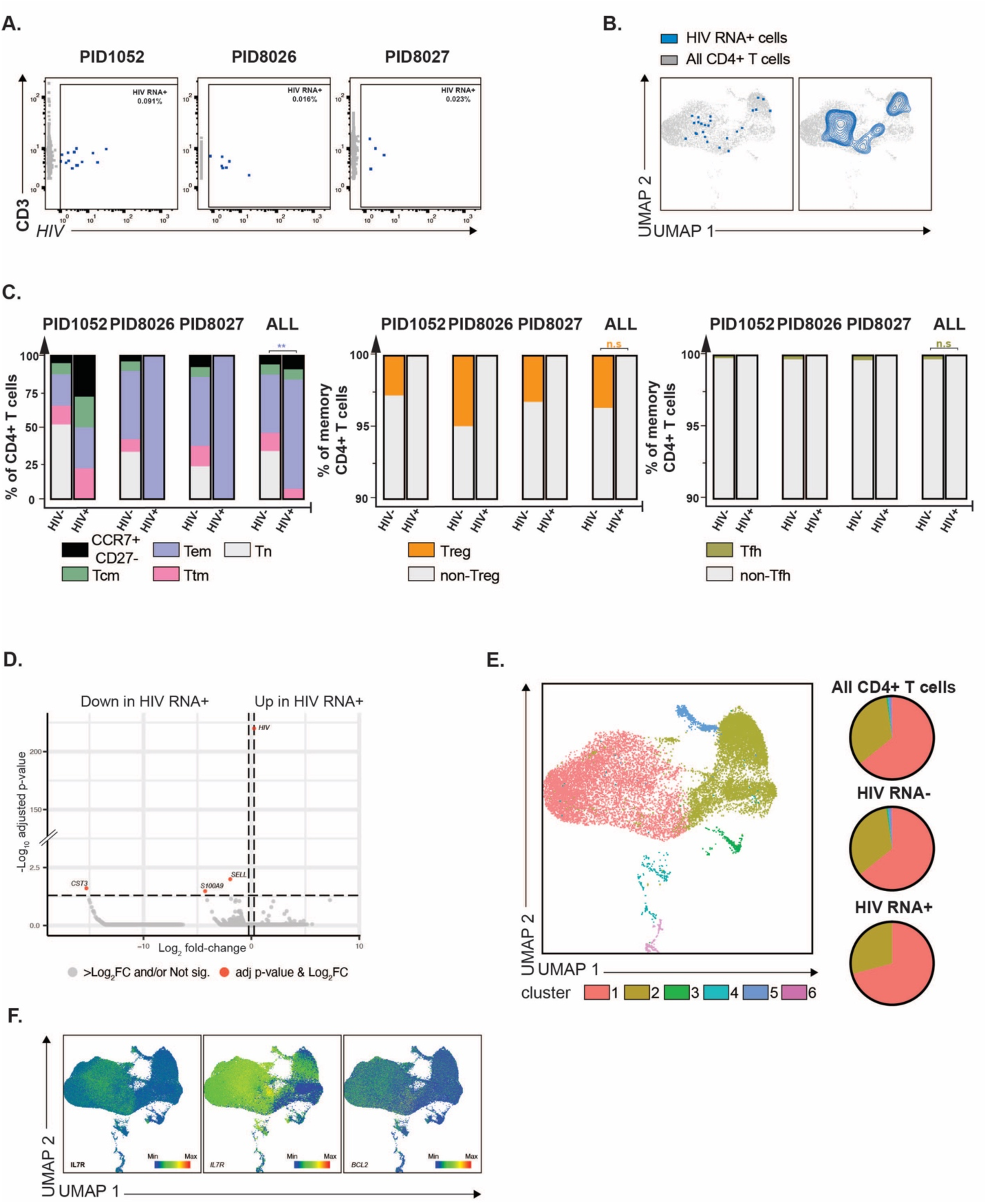
Most HIV RNA+ cells from ART-suppressed PWH exhibit stem-like rather than cytolytic features. **A.** CD3+CD4+ T cells expressing HIV RNA were identified by HIV-seq from 3 ART-suppressed PWH (PID1052, PID8026, and PID8027) based on HIV RNA levels. HIV RNA+ cells are depicted in blue and HIV RNA- cells are depicted in gray. Percentages of HIV RNA+ cells are indicated in the upper right of each plot. **B.** A diverse array of HIV RNA+ cells exists among CD4+ T cells from ART-suppressed PWH. Shown are UMAP plots depicting HIV RNA+ cells as blue dots (*left*) or contours (*right*), against a background of HIV RNA- CD4+ T cells depicted in gray, for all 3 donors listed in *panel A* combined. **C.** HIV RNA+ cells from ART-suppressed PWH are enriched among Tem cells. Shown in the top panel are bar graphs depicting distribution of Tn, Tcm, Tem, Ttm, memory CCR7+CD27- cells among HIV RNA- and HIV RNA+ CD4+ T cells. The remaining panels depict the proportion of Treg and non-Tregs, and Tfh and non-Tfh, among HIV RNA- and HIV RNA+ memory CD4+ T cells. **P ≤ 0.01, ***P≤ 0.001, ****P≤ 0.0001 as determined by generalized linear mixed model (GLMM). **D.** HIV RNA+ cells from ART- suppressed PWH express low levels of Tcm marker *SELL*, alarmin *S100A9,* and cysteine protease inhibitor *CST3*. Shown is a volcano plot displaying differentially expressed transcripts in HIV RNA+ vs. HIV RNA- CD4+ T cells, with select transcripts annotated. Red dots correspond to transcripts with 0.25log_2_ fold-change expression and with adjusted p value ≤ 0.05, as determined by genewise quasi F-tests. **E.** HIV RNA+ cells are not enriched among cytolytic cluster 2. *Left:* UMAP depicting the 6 clusters of CD4+ T cells identified by Louvain clustering. *Right:* Pie graphs showing the distribution of total CD4+ T cells (*top*), HIV RNA- CD4+ T cells (*middle*) or HIV RNA+ CD4+ T cells (*bottom*) among the different clusters. **F.** Cluster 1, which contains the biggest proportion of HIV RNA+ cells, is enriched for *IL7R* and *BCL2* transcripts, as well as IL7R protein, which are characteristic of long-lived and stem-like T cells. Heatmaps depict relative expression of the indicated transcripts and protein.

Louvain clustering analysis revealed that unlike HIV RNA+ cells collected during viremia, those collected during suppression were not enriched among cytolytic cluster 2. Instead, the cluster distribution of the infected cells mirrored that of the overall CD4+ T cell population (Fig. 4E). Hence, the HIV RNA+ cells were primarily distributed among cluster 1, the most abundant cluster. Cluster 1 primarily consists of memory CD4+ T cells expressing high levels of IL7R, the α chain of the IL7 receptor (also known as CD127), suggesting they are long-lived cells with stem-like proliferative capacity (Fig. 4F). Further supporting the notion of their being long-lived memory cells is their preferential expression of *BCL-2* (Fig. 4F), a pro-survival, anti-apoptotic gene implicated in HIV persistence^45–48^. Taken together, these findings suggest that in blood, the majority of HIV RNA+ cells during ART suppression reside not within cytolytic CD4+ T cells – as was observed during viremia – but rather within a long-lived population of memory CD4+ T cells.

### CD4+ T cells exhibit stronger antiviral immunity and upregulate the integrated stress response pathway during viremia as compared to after ART suppression

Leveraging the fact that HIV-seq was performed on paired specimens from before vs. after ART suppression, we then compared the transcriptomes of total CD4+ T cells across these two conditions. Total CD4+ T cells coming from viremic vs. suppressed specimens were transcriptionally divergent, as reflected by distinct UMAP localizations (Fig. 5A, Fig. S6), even though their distribution among classical CD4+ T cell subsets was similar (Fig. 5B). Although DEGs were identified (Fig. 5C, Table S5), none retained statistical significance following correction for multiple comparisons. As an exploratory analysis, however, we assessed the DEGs with the lowest raw p-values to gain insights into potential cellular pathways distinguishing CD4+ T cells during active viremia from those following ART suppression. This revealed CD4+ T cells during viremia to preferentially express higher levels of cytotoxic and proinflammatory genes (*SP100, IL6*), and genes related to interferon α, β and γ signaling (*IFI27, IFI44L, IFIT3, IFI6, IFI44, ISG15, IFITM10, IFI30, IFI27L1, MX1, IFIH1, IFNLR1, IFIT5, NEAT1, TYMP*). Some of these (*IFI44L, ISG15, NEAT1, TYMP)* have previously been reported to be higher among CD4+ T cells during active viremia as compared to ART treatment^49^. This upregulation may reflect an antiviral host response triggered by high levels of viral transcripts and proteins during viremia. Other highly expressed genes during viremia included cytolytic genes (*GZMB, GZMK*) whose expression can be induced by cytokines such as IL2 and IL15^50^, and during inflammation and viral infections^51^.

**Figure 5.**
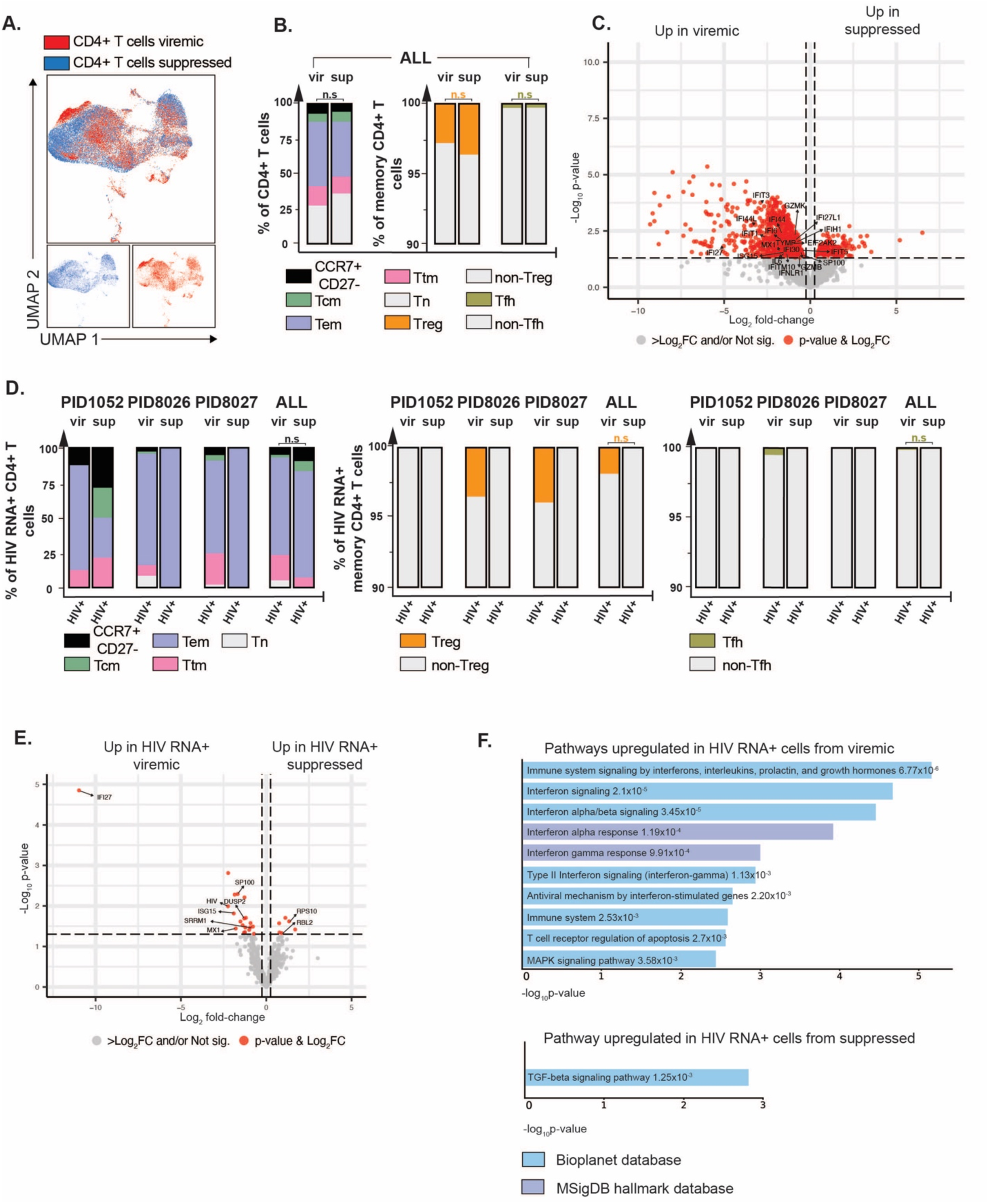
HIV RNA+ cells from viremic PWH exhibit a pro-inflammatory and anti-viral state while those from ART-suppressed PWH exhibit properties that favor senescence and HIV restriction. **A.** ART suppression elicits global changes in host transcriptome. Shown are UMAP plots of total CD4+ T cells from viremic (red) vs. suppressed (blue) timepoints from PID1052, PID8026 and PID8027 combined. **B.** Distribution among classic CD4+ T cell subsets is not altered during ART suppression. Shown are bar graphs depicting distribution of Tn, Tcm, Tem, Ttm, memory CCR7+CD27- cells as well as the distribution of Treg, non-Treg, Tfh, and non-Tfh among total and memory CD4+ T cells as indicated during viremia (vir) and upon ART suppression (sup). Results from all 3 donors are combined. n.s. as determined by generalized linear mixed model (GLMM). **C.** Compared to CD4+ T cells during ART, CD4+ T cells during viremia express higher levels of host transcripts involved in immune responses and cytotoxicity. Shown is a volcano plot displaying differentially expressed transcripts in CD4+ T cells from viremic vs. suppressed time points, with select transcripts annotated. Red dots correspond to transcripts with 0.25log_2_ fold-change expression and with p value ≤ 0.05, as determined by gene-wise quasi F-tests. **D.** Distribution of HIV RNA+ cells among classic CD4+ T cell subsets is not altered during ART suppression. Shown are bar graphs depicting distribution of Tn, Tcm, Tem, Ttm, memory CCR7+CD27- cells as well as the distribution of Treg, non-Treg, Tfh, and non-Tfh among HIV RNA+ memory CD4+ T cells, during viremia (vir) and upon ART suppression (sup). Results from all 3 donors are combined. n.s. as determined by generalized linear mixed model (GLMM). **E.** HIV RNA+ cells during viremia express high levels of Interferon Stimulated Genes (ISGs) and low levels of transcripts that restrict HIV. Shown is a volcano plot displaying up- and down- regulated transcripts in HIV RNA+ cells during viremia vs. ART suppression, with select transcripts annotated. Red dots correspond to transcripts with 0.25log_2_ fold-change expression and with a non-adjusted p value ≤ 0.05, as determined by gene-wise quasi F-tests. **F.** HIV RNA+ cells preferentially exhibit gene expression signatures of interferon signaling, antiviral and immune responses during viremia, and of TGF-β signaling during ART suppression. Shown is pathway analysis comparing HIV RNA+ cells in the context of viremia vs. suppression, using the Bioplanet 2019 and MSigDB hallmark databases. The p-value representing the statistical significance of enrichment of the gene set withing the pathway is indicated next to each pathway.

Also of interest was that that during viremia, CD4+ T cells increased expression of *EIF2AK2* (Fig. 5C, Table S5), a key gene involved in integrated stress response (ISR), a pathway previously reported to be induced during acute HIV infection^52,53^. During ISR, *EIF2AK2* expression is induced by T1IFNs, which then upon binding to viral dsRNA can initiate a cascade of events culminating in diminished protein translation. Indeed, when we performed DEG analysis of all donors combined using an approach previously implemented to identify DEGs between total CD4+ T cells during active HIV viremia vs. after ART suppression^25,49^ (see Methods), we observed downregulation of numerous ribosomal transcripts (RPL and RPS transcripts) during viremia (Fig. S7 and Table S6). The diminished ribosomal transcript levels were accompanied by diminished expression of *EEF2*, whose downregulation has been shown to reduce active ribosomes^53^ and which shuts down mRNA translation resulting in overall diminished viral protein synthesis^54^. Hence, the ISR pathway, which serves to coordinate cellular responses to various stressors by regulating protein synthesis and gene expression and is a host response to limit viral spread, appears to be a characteristic feature of acute untreated HIV infection that is turned off upon ART suppression.

### Relative to HIV RNA+ cells during viremia, those during ART suppression upregulate TGFβ signaling pathways and exhibit diminished activation of host responses

We then compared the HIV RNA+ CD4+ T cells from the paired specimens. Classical CD4+ T cell subset distributions of HIV RNA+ cells between the two timepoints were similar, with preferential distribution among Tem cells for both (Fig. 5D). Here again, multiple DEGs were identified between the timepoints, but none retained statistical significance after correction for multiple comparisons. As an exploratory analysis, we assessed the identities of the DEGs based on the raw p-values (Fig. 5E and Table S7). This analysis revealed upregulation of multiple ISGs and proinflammatory genes in HIV RNA+ cells during viremia compared to suppression, including *IFI27*, *MX1*, *ISG15*, *DUSP2,* and *SP100*. These genes were also upregulated among total CD4+ T cells during viremia (Fig 5C and Table S5). IFI27 in particular was increased by more than 10-fold in HIV RNA+ CD4+ T cells during viremia as compared to those during ART. Given that HIV-1 Vpr and Tat can directly induce *IFI27* production^55–57^, and that expression of these viral proteins may be diminished in the context of ART suppression due to multiple blocks to HIV transcription^7–9^, it is possible that elevated expression of Vpr and Tat is driving *IFI27* expression prior to ART initiation. Interestingly, *IFI27* levels correlate with inflammation and disease progression during both HIV-1 and HIV-2 infection^58,59^, and may contribute to HIV pathogenesis^58,59^.

DEGs associated with HIV transcription were also upregulated in HIV RNA+ cells during viremia as compared to after ART suppression. HIV RNA+ cells during viremia expressed higher levels of *SRRM1*, a modulator of HIV-1 splicing that is involved in the regulation of Tat and Nef expression^60^, and lower levels of two genes implicated in silencing HIV transcription and translation: *RBL2*, which recruits and targets histone methyltransferases, leading to epigenetic transcriptional repression; and *RPS10*, which binds HIV Nef to form a complex that decreases viral protein synthesis^61^. Together, these findings suggest that during viremia, HIV RNA+ cells exhibit a transcriptional profile that favors the production of more HIV transcripts. This finding accords with our having observed elevated HIV transcript levels among HIV RNA+ cells during viremia as compared to during suppression in two out of our three participants (11- and 14-fold increase, respectively, for PID8026 and PID8027).

Pathway analysis of the DEGs in HIV RNA+ cells before versus after ART suppression supported the notion that HIV RNA+ cells during viremia are preferentially in an activated, antiviral state, characterized by upregulation of multiple interferon pathways (IFN I and IFN II) (Fig. 5F, top). Conversely, HIV RNA+ cells during ART suppression preferentially upregulated TGF-β signaling (Fig. 5F, bottom). This entailed upregulation of *RBL2* and *ITGB1* (Table S5), genes associated with the TGF-β signaling pathway. *ITGB1*- and *RBL2*-associated TGF-β activation has been implicated in tumor suppression and cancer growth arrest^62,63^. Although *ITGB1* and *RBL2* have not been directly implicated in HIV infection, TGF-β signaling has been recently implicated in promoting HIV latency^64,65^. Our data suggest that this TGF-β-associated signature is a phenomenon that only emerges in the context of ART suppression, as HIV RNA+ cells during viremia do not exhibit such a signature. Overall, these results indicate that the transcriptional profiles of HIV RNA+ cells differ depending on whether or not ART is present, and that the features of HIV RNA+ cells during viremia cannot be assumed to be the same as those during ART suppression.

## DISCUSSION

In this study, we introduce HIV-seq as a method to improve the efficiency by which individual HIV RNA+ cells can be identified by scRNA-seq. Applying HIV-seq to blood samples collected at viremic and suppressed timepoints from the same set of individuals allowed us to discover features of infected cells in viremic individuals, and discern differences between these cells and active reservoir cells that persist during stable ART suppression.

HIV-seq increased detection of HIV RNA+ cells from PWH by up to 44%, and enabled in-depth scRNA-seq of the highest numbers of HIV RNA+ cells reported to date, in the context of both viremia and ART suppression. We recovered and analyzed 1,072 HIV RNA+ cells from four viremic donors, in comparison to prior methods using classic scRNA-seq^32^, ECCITEseq^25^, and DOGMAseq^49^, which had identified 164 total cells from 14 donors (with one highly viremic donor further sequenced to gain an additional 223 HIV RNA+ cells), 81 total cells from 6 donors, and 256 total cells from 6 donors, respectively. Our identification of 26 HIV RNA+ cells from 3 donors on ART is also substantially higher than prior studies, which had reported 2 total cells from 14 donors^32^, 9 total cells from 6 donors^25^, and 14 total cells from 6 donors^49^.

The elevated numbers of HIV reads identified by HIV-seq also enabled an in-depth analysis of the distribution HIV transcripts among infected cells. We found reads spanning the entire HIV genome, regardless of whether we included HIV capture oligos. During viremia, the most frequently detected transcript was *pol*, followed by *gag*, *vif*, *env*, and *tat*. Likewise, *gag* and *pol* were most common during ART suppression. This distribution likely reflects three key factors. First, the technology used for scRNA-seq can introduce biases at the stages of RNA capture (use of poly-dT +/- specific sequences), reverse transcription, binding of template switch oligo (to CCC trinucleotides), second strand synthesis, amplification (amplicon size), fragmentation, ligation, sequencing, and analysis (location of cell/transcript barcodes). Second, alignment efficiency is impacted by the extent to which a sequenced read matches our subtype B consensus sequence. *Gag* and *pol* genes are the most conserved among subtype B HIV-1 isolates^66,67^, which may have contributed to their being the most frequently identified HIV transcripts in our scRNA-seq analysis. Lastly, read distribution can be affected by the relative proportions of each transcript type. In particular, blocks to HIV transcription in active reservoir cells primarily occur before the *env/nef* regions^7^, resulting in a higher presence of 5’ reads in the samples. All these factors together may have accounted for our preferential identification of *gag* and *pol* transcripts, irrespective of HIV-seq.

We identified over a thousand HIV RNA+ cells by HIV-seq in the absence of ART, enabling an in-depth analysis of the features of HIV-infected cells during viremia. This analysis both confirmed prior studies analyzing fewer cells and also revealed new insights not previously reported. HIV RNA+ cells during viremia were predominantly of the Tem phenotype, consistent with previous reports^22,25^. One likely explanation is the increased susceptibility of Tem cells to HIV infection relative to their Tcm counterparts^24^. Also consistent with prior studies^25,32,49^ is our observation that HIV RNA+ cells preferentially reside in a cluster of cells exhibiting a Th1/cytotoxic phenotype, defined by preferential expression of Th1-defining markers along with cytolytic markers including granzymes, perforin, and granulysin. Active viral replication may promote the acquisition and maintenance of cytotoxic functions by CD4+ T cells by eliciting a sustained pro-inflammatory environment, which has been shown in the context of cancer^68^ as well as during influenza infection^69^.

We also discovered that during active viremia, HIV RNA+ cells expressed lower levels of the restriction factors *APOBEC3A* and *SERPINA1* compared to their HIV RNA- counterparts, which to our knowledge has not previously been reported and may reflect an immune evasion mechanism mediated by the virus. APOBEC3A is a DNA cytidine deaminase that exhibits antiviral activity, including against HIV^70,71^, and can also help maintain HIV latency in CD4+ T cells through recruitment of epigenetic silencing machinery to the LTR^72^. The HIV accessory gene *Vif*, however, can target APOBEC3A for proteasome-mediated degradation^73^. Our finding of decreased *APOBEC3A* expression at the transcript level suggests that there may be mechanisms beyond proteasome-mediated degradation to suppress APOBEC3A activity.

SERPINA1 is a restriction factor that is induced during inflammation and inhibits HIV LTR-driven transcription^74^, and whose expression can be regulated by methylation, independently of inflammation^75^. The extent to which HIV RNA+ cells downregulate SERPINA1 expression through methylation is unknown, but this mechanism is conceivable given the profound epigenetic changes induced by HIV infection^76^. Hence, diminished expression of both *APOBEC3A* and *SERPINA1* in HIV RNA+ cells during acute viremia may promote rapid production of new virions by promoting HIV transcription.

In addition to analyzing HIV RNA+ cells, we also leveraged our in-depth sequencing datasets to assess the extent to which active HIV viremia affects the transcriptomes of total CD4+ T cells. Comparing the scRNA-seq profiles of total CD4+ T cells before and after ART suppression revealed an upregulation of proinflammatory genes and genes related to interferon α, β, and γ signaling during viremia. This finding aligns with a recent demonstration of elevated type I IFN gene expression (*IFI44L, ISG15, XAF1, NEAT1, TYMP, TRIM22*) in total CD4+ T cells during untreated HIV infection^49^, and is consistent with the notion of a pro-inflammatory response induced by active viral replication.

Unexpectedly, we also discovered that total CD4+ T cells during viremia upregulated the ISR pathway relative to their counterparts during ART suppression. In general, ISR serves as a protective host response against viruses by reducing global protein synthesis to inhibit viral replication, and in some cases can further induce apoptosis of infected cells^52^. However, some viruses – including HIV – seem to have evolved mechanisms to hijack or benefit from ISR^52,77^.

For example, the ISR-associated transcription factor ATF4 can bind to the HIV promoter to stimulate HIV transcription^77^. It is thus possible that global upregulation of ISR among CD4+ T cells creates an intracellular environment favorable for HIV gene expression, thereby facilitating rapid systemic spread of the virus during untreated infection.

In addition to revealing insights into HIV pathogenesis and spread during untreated infection, HIV-seq also enabled in-depth analysis of active reservoir cells in aviremic individuals. To date, no study has conducted a comprehensive scRNA-seq analysis specifically on HIV RNA+ cells in the context of ART suppression, as prior studies have either combined HIV RNA+ cells from viremic and suppressed samples for analysis (due to low numbers of HIV RNA+ cells identified during ART)^25,32^ or only performed primary analysis on HIV RNA+ cells during viremia^49^. Importantly but perhaps not surprisingly, we found that HIV RNA+ cells during ART suppression differ from those during active viremia, suggesting that HIV RNA+ cells during viremia should not be used as a proxy for understanding how HIV persists in ART-suppressed PWH. Although HIV RNA+ cells in both instances were predominantly Tem, those during ART suppression did *not* preferentially harbor a cytotoxic phenotype. This finding can be explained by the general short-lived nature of effector/cytotoxic lymphocytes, which has been described for CD8+ CTLs^78^. By contrast, HIV reservoir cells are long-lived, and recent studies of total and genome-intact HIV reservoir cells have suggested preferential expression of markers of cell survival ^79,80^. Although it may seem perplexing that we observed active reservoir cells to preferentially reside among Tem, which are generally considered short-lived^81^, it is worth noting that long-lived Tem cells have been described in the context of viral infections^82–84^.

Supporting the notion that active reservoir cells, like the total reservoir cells, exhibit features of longevity, we found that on-ART HIV RNA+ cells preferentially resided in a cluster of cells expressing high levels of CD127, the alpha chain of the IL7 receptor, which is a major driver of homeostatic proliferation. IL7 can promote stabilization of a long-lived reservoir of HIV- infected cells^85^, and is associated with a slower contraction of the total HIV reservoir (as defined by HIV DNA levels) over time^86^. Our data suggest that IL7 may also drive persistence of the active reservoir. We also found this dominant cluster of HIV RNA+ cells to preferentially express BCL-2, an anti-apoptotic protein that inhibits apoptosis by regulating mitochondrial membrane permeability and preventing the release of pro-apoptotic factors^87^. Interestingly, *ex vivo* treatment of cells from ART-suppressed PWH with different BCL-2 inhibitors such as venetoclax^45^ or obatoclax^48^ decreases the pool of genome-intact HIV-infected cells.

Furthermore, venetoclax delays viral rebound upon ART interruption in a humanized mouse model of HIV persistence^45^. Our data suggest that the active reservoir, like genome-intact reservoir cells, should also be sensitive to BCL-2 inhibitors. The outcome of a recently initiated clinical trial testing venetoclax as a therapeutic approach to achieve HIV remission in ART- suppressed PWH ^88^ will be interesting in that regard.

Besides exhibiting stem-like properties, active reservoir cells also exhibited increased activation of the TGF-β pathway. Recent studies in a non-human primate model of SIV infection implicated TGF-β in promoting HIV persistence, through mechanisms related to both the cytokine’s immunosuppressive effects as well as its ability to suppress viral gene expression^64,65^. Our finding that active reservoir cells preferentially activate the TGF-β pathway – in particular by upregulating ITGB1, which mediates release of the active form of TGF-β^62^, and RBL2, which in response to TGF-β activation mediates changes cell cycle progression^63^ – suggests that persistent HIV may utilize the TGF-β pathway to achieve immune evasion.

Immune evasion may be particularly important for active reservoir cells, as these cells can express HIV proteins that can then be processed for recognition by HIV-specific CD8+ T cells^89–91^. Our finding that active reservoir cells utilize the TGF-β pathway, alongside evidence that galunisertib (a TGF-β1 receptor inhibitor) promotes *ex vivo* reactivation of HIV from cells of ART-suppressed PWH^64^, favors the notion that targeting the TGF-β pathway may be a viable approach for the “kick and kill” strategy for eliminating HIV reservoir cells.

Together, these results demonstrate that HIV RNA+ cells during active viremia are primarily cytotoxic CD4+ T cells with diminished expression of restriction factors targeting HIV transcription, while those during ART suppression exhibit features enabling long-term survival through anti-apoptotic and homeostatic proliferation mechanisms along with exploitation of TGF- β signaling pathways to achieve immune evasion. Future studies should apply HIV-seq to more broadly characterize the active reservoir, for example in the context of tissues where HIV primarily persists. In addition, applying HIV-seq in the context of clonal expansion analysis using single-cell VDJ analysis, and further developing this technique to allow for multiplexing with other platforms (e.g. single-cell ATACseq) will have utility in furthering our understanding of the mechanisms by which HIV can persist long-term despite ART suppression of viremia.

## METHODS

### Ethics statement

The study was approved by the Committee on Human Research (CHR), the Institutional Review Board for the University of California, San Francisco (approval #11–07551 and #10- 01561). All study participants provided written informed consent.

### Study Population

The study participants were HIV-infected adults on suppressive ART (median age = 38.5; median CD4 count = 490 cells/mm^3^; Table 1). Paired and longitudinal, archived PBMC samples were obtained from the UCSF Treat Acute Study and the SCOPE cohort. Samples were collected prior to ART initiation (Week 0) and following ART suppression (Week 24: PID8026; Week 45: PID8027; and Week 70: PID1052). A total of two aliquots of 10^7^ cells each were obtained for each time point (viremic and suppressed), with one aliquot preserved for HIV DNA/RNA measurements. An additional participant (PID0145), for whom only the viremic (Week 0) time point was available, was recruited from the San Francisco VA Medical Center.

**Table 1.**
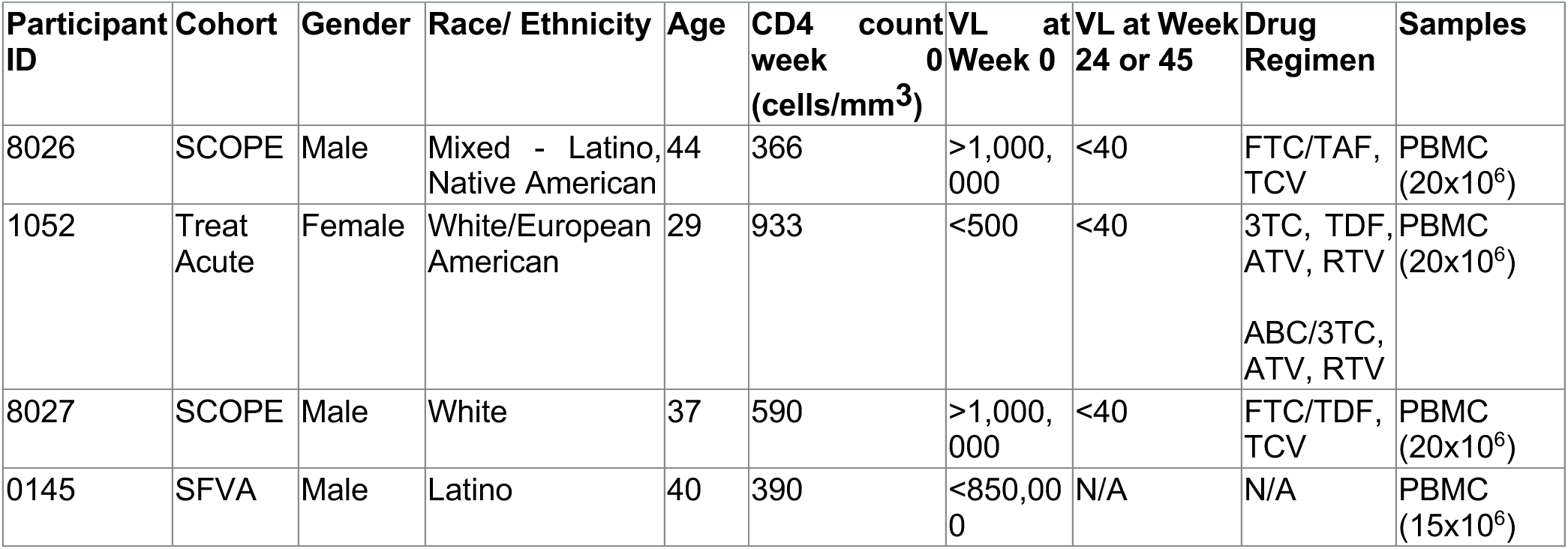
Characteristics of study participants living with HIV.

### HIV DNA and RNA measurements

For each participant, levels of HIV DNA and RNA were evaluated to ascertain whether their reservoir was sufficiently high for detection with limited cell inputs (up to 10,000 cells per well). An aliquot of 10^7^ cryopreserved cells was tested from each time point (viremic and suppressed) for each donor. Each aliquot was thawed, cryopreservation medium was removed, and total RNA and DNA were extracted using TRI Reagent (Molecular Research Center, Inc., Cincinnati, OH) as per manufacturer’s instructions, with the following modifications: polyacryl carrier (Molecular Research Center, Inc., Cincinnati, OH) was added to TRI reagent prior to lysis, RNA was resuspended in RNase free-water, DNA was extracted using back extraction buffer (4M guanidine thiocyanate, 50mM sodium citrate, 1M Tris), polyacryl carrier was added to the aqueous phase containing the DNA, and DNA was resuspended in QIAGEN buffer EB^8^.

Reverse transcription and droplet digital PCR were performed as previously described^7^. HIV DNA (R-U5-pre-Gag region) and copies of the housekeeping gene TERT (telomere reverse transcriptase) were measured in duplicate by droplet digital (dd)PCR (Bio-Rad QX100). A threshold of 3 HIV DNA copies/10,000 cells at the suppressed time point was set as a threshold for study inclusion, as such levels yielded a reasonable likelihood of detecting HIV transcripts at the single cell level using the 10X Genomics scRNA-seq platform. All samples subjected to single-cell sequencing in this study met this threshold.

### scRNA-seq sample preparation

#### PBMC

A total of 10^7^ PBMCs were thawed at 37°C and washed once with warm media (RPMI [Corning Inc., Corning, NY] supplemented with 10% FBS [VWR, Radnor, PA]) and then resuspended in FACS buffer (RPMI, supplemented with 2% FBS and 2 mM EDTA [Thermo Fisher Scientific]) prior to counting. After setting aside 10^5^ cells from each sample for downstream PBMC spike-in, CD4+ T cells were purified from the remaining cells by negative selection using the EasySep Human CD4 T cell enrichment kit (StemCell). The PBMCs from the same donor that were set aside were then spiked back in at a ratio of 5:100 in order have non- CD4+ T cells to help establish gates for defining HIV RNA+ cells.

#### TotalSeq-C Antibody Staining

TotalSeq-C pooled antibody mix (Biolegend) was prepared so as to contain 0.4 μg of each antibody in a final volume of 100 μL/sample, in PBS (Ca++ and Mg+ free, Corning Inc.) containing 3% FBS, as per devised panels (Table S1). A million cells were pelleted, resuspended in RPMI supplemented with 3% FBS, and incubated at 4°C for 10 min with Fc block (Miltenyi) at a 1:10 dilution. 100 μL/sample of TotalSeq pooled antibody mix was then added to the cells, which were incubated for 30 min at 4°C. Cells were then washed three times in RPMI containing 0.04% BSA, strained using a 40 μm cell filter (BD Falcon), and resuspended at a concentration of 1000 cells/μL.

### Custom HIV primers

Custom HIV-seq primers were designed by appending HIV-specific capture sequences (Table S2) to a non-poly(dT) PCR handle (Fig. 1). The concentration of 10X Poly(dT) oligo (poly-dT RT Primer PN 2000007) is estimated to be 1.33 μM in each 10X reverse transcription (RT) reaction^92^. Initial experiments spiking in different concentrations of PreGag primers revealed that 0.67 μM of primer resulted in lower UMIs mapped across the HIV genome relative to 0.33 μM primer. Furthermore, the 0.67 μM concentration resulted in a lower frequency of detected HIV transcripts. When we iteratively lowered primer concentrations, we found the optimal HIV primer concentration to be 41.6 nM. The primer pool (containing 41.6 nM of each primer) was added to the 10X Genomics reverse transcriptase (RT) reaction mix containing RT Reagent B, Poly-dT RT Primer, Reducing Agent B, and RT Enzyme C (HIV-Seq RT Mix), and used for reverse transcription of encapsulated cells as described below.

### Reverse transcription, library preparation and sequencing

Chromium Next GEM Single Cell 5’ Reagent Kits v2 (Dual Index) with Feature Barcode technology for Cell Surface Protein & Immune Receptor Mapping (10X Genomics, PN1000263) and the Chromium^TM^ Controller (10X Genomics, PN120223) were used for gene expression library and cell-surface protein library generation. Briefly, 20 μl of a TotalSeqC-stained single-cell suspension (∼1000 cells/µl) was mixed with the HIV-Seq RT Mix, barcoded using Single Cell VDJ 5’ Gel Beads, and partitioned with oil onto a Chromium Next GEM Chip K. The chip was then loaded onto a Chromium Controller for single-cell GEM generation and barcoding. Reverse transcription reactions were performed according to the Chromium Single Cell 5’ Reagent Kits v2 (Dual Index) User Guide (10x Genomics CG000330, Rev A). Sequencing libraries were constructed with 13 cycles of PCR during cDNA amplification and 14 cycles of Sample Index PCR. Gene expression and cell surface protein libraries were pooled and sequenced on a the NovaSeq 6000 lane (S4 flow cell) and sequenced at a minimum of 50,000 reads / cell.

### Data processing, statistics, and analysis

Sequencing libraries from n=4 independent experiments were analyzed. A custom Human/HIV consensus subtype B reference sequence was purpose-built for this study by downloading a Group M alignment from the Los Alamos National Laboratory’s HIV sequence database (2018 version), filtering for subtype B sequences, realigning using MAFFT^93^ with default settings, and generating a majority consensus sequence using Geneious (http://www.geneious.com/). The consensus HIV sequence was appended to the human reference genome (GRCh38-2020) and annotated as a single exon. Alignment of scRNA-seq reads, collapsing of reads to unique molecular identifier (UMI) counts, and cell calling was performed using CellRanger 6.0.2 (10X Genomics). Filtered count matrices of features generated using the CellRanger count function were then subjected to multiple cleanup steps. Cells with high mitochondrial genes expression (mtDNA% > 15%) and low features (<500) were removed, along with doublet cells (identified using the DoubletFinder package^94^), in R. All samples were normalized for their transcriptome library depth and batch-corrected between donors using the Seurat^95^ NormalizeData and integration function from Seurat package, in R. Downstream analysis was performed in SeqGeq (mostly for visualization) (FlowJo, LLC) and Seurat v4.3.0. In all analyses, genes are depicted in italics and proteins in bold.

#### Clustering

Graph-based clustering was performed using the Louvain algorithm implementation^96^ in the FindClusters Seurat function. The optimal clustering resolution parameters were determined using Random Forests^97^ and a silhouette score-based assessment of clustering validity and subject-wise cross-validation^98^. T cell Receptor Alpha Variable (*TRAV*) and T cell Receptor Beta Variable (*TRBV*) genes were removed from the variable features used for clustering as they were driving the clustering in a donor-specific manner, as to be expected with randomly-generated VDJ sequences. Six distinct biologically relevant clusters (clusters 1–6) were identified, which were used for further analyses.

#### Manual gating

Manual gating was conducted using SeqGeq software to delineate classic CD4+ T cell subsets and to identify HIV-expressing cells. Both gene and protein expressions were used for gating.

#### Statistical Analysis of CD4+ T Cell Subsets and HIV RNA+ Cell Distribution

For establishing associations between CD4+ T-cell subset proportions among the HIV RNA- and HIV RNA+ cells, and HIV RNA+ cell proportions among the different clusters, a generalized linear mixed model (GLMM) with a binomial probability distribution implemented in the lme4^99^ package in R was used. In the model, CD4 subset or cluster membership was treated as the outcome being studied. This was represented by comparing the number of cells within the subset/cluster to the number of cells outside of it. The change in subset/cluster membership between HIV RNA status and timing of measurements (viremic vs suppressed) was estimated as a log odds ratio, defined as the change in the log odds of subset/cluster membership. This was estimated with the emmeans^100^ R package using the GLMM model fit.

#### Differential expressed gene (DEG) analysis

Pseudobulk DEG analysis was performed to identify genes or proteins that were significantly upregulated or downregulated between different cell populations or experimental conditions. DEG analysis at the RNA level was carried out using the R package muscat^101^, and counts were summed across clusters and samples. The pbDS function to estimate associations with disease, was run with the edgeR^102^method, minimum cells set to 3, filter set to gene, and donor was included as a confounder in the model. In the exploratory analysis comparing total CD4+ T cells during viremia vs. suppression, all donors within a group (viremic or ART- suppressed) were combined, and then gene expression levels were compared across individual cells within each group, followed by implementation of the Wilcoxon Rank-sum test to assess for DEGs and proteins, similar to analytical approaches recently described^25,49^. Of note, this analytical approach should be considered exploratory as it does not fully account for correlations between cells from the same subject or paired study designs^103,104^. Results visualized as volcano plots were plotted using EnhancedVolcano^105^. Select genes and proteins of interest among parameters that passed the indicated adjusted p-value (or raw p-value where indicated) thresholds of 0.05 and a log_2_ fold-change greater than 0.25 are highlighted in the volcano plots. All genes are depicted in italics, and proteins in bold. T cell receptor (TCR) genes (*TRAV* and *TRBV*) were deliberately omitted from the analysis, since private TCR sequences drove donor-specific effects, and the primary focus of our study was to identify gene expression profiles shared across donors.

#### Pathway Enrichment Analysis

The differential gene expression analysis results filtered by adjusted p-value < 0.05 and log_2_ fold-change > 0.25 were subjected to over-representation analysis (ORA) using the Enrichr^106–108^ web-based tool. Pathway analysis was performed against two curated databases: BioPlanet 2019 and MSigDB Hallmark 2020. This identified significantly enriched pathways and functional categories along with statistical metrics and p-values.

### Distribution of HIV reads

An additional analysis assessing distribution of reads across the HIV genome was performed to determine where HIV transcripts most frequently align. We selected PID8027 and PID1052, for which we had data on the absence versus presence of HIV-capture oligos. Gene expression reads from the week 0 (viremic) timepoint were aligned to a custom, annotated HXB2 reference sequence, which defined discrete HIV coding regions (‘5’LTR U3’, ‘5’LTR R’, ‘5’LTR U5’, ‘Start of Gag’, ‘Gag P1’, ‘Pol’, ‘Vif’, ‘Vpr’, ‘Rev’, ‘Vpu’, ‘Env’, ‘Tat’, ‘Nef’, ‘3’ LTR U3’, ‘3’LTR R’) using CellRanger (10X Genomics, version 6.0.2). The distribution of HIV reads aligning to these specific regions was then filtered, quantitated using an R script and compared between samples subjected to the conventional 10X Genomics’ 5’ scRNA-seq pipeline vs. HIV- Seq.

## Acknowledgments

This work was supported by the National Institutes of Health (P01AI169606, R01DK120387, R01AI132128, R01AI147777, R01DK131526, R01AI183286, R21AI170166, UM1AI164559, UM1AI164567, UM1AI164560) and the California HIV/AIDS Research Program ([S.T] BB19-SF-009/A135087). We also acknowledge support from CFAR (P30AI027763) and the James B. Pendleton Foundation. S.T is supported by Doherty Institute for Infection and Immunity Locarnini Fellowship in Virology and University of Melbourne Department of Infectious Diseases Research Support Package. The funders had no role in study design, data collection and analysis, decision to publish, or preparation of the manuscript. We thank Viva Tai and Marian Kerbleski for assistance with the SCOPE specimens; Vivian Pae, Sannidhi Sarvadhavabhatla, and Maria Sophia Donaire for assistance with the Treat Acute HIV specimens, Francoise Chanut for editorial assistance; and Robins Givens for administrative assistance; and the study participants for their samples. This work utilized the computational resources of the UCSF Wynton cluster (https://wynton.ucsf.edu).

## Author Contributions

J.F and S.T. designed the experiments, performed scRNA-seq experiments, conducted analyses, interpreted data, and prepared figures and tables. X.L. developed pipeline for quality control analysis of scRNA-seq datasets. N.E., R.T., and D.A. performed scRNA-seq analyses, and P.R. created the HIV consensus genome used for scRNA-seq data alignment. R.H. and J.K.W. recruited participants, collected clinical data, and collected biospecimens. S.G.D. oversaw the SCOPE cohort procedures and S.A.L. managed specimen collection. A.J.B., N.R.R. and S.Y. performed supervision. J.F, S.T, N.R.R., and S.Y. conceived the study, interpreted data, and wrote the manuscript. All authors have read and approved this manuscript.

