## Supplementary Figures and Tables for "HIV-SEQ Reveals Global Host Gene Expression Differences Between HIV-Transcribing Cells from Viremic and Suppressed People with HIV"

### SUPPLEMENTARY TABLES

**Table S1.** TotalSeq C antibody panel for CITE-seq analysis

| ID | Name | Read | Pattern | Sequence | Feature_type | Supplier ID | Clone | Other Names | Isotype |
|---|---|---|---|---|---|---|---|---|---|
| CD127 | CD127_TotalC | R2 | 5PNNNNNNNNNN(BC) | GTGTGTTGTCCTATG | Antibody Capture | TotalSeq™-C0390 anti-human CD127 (IL-7Rα) Antibody | A019D5 | IL-7 receptor α chain, IL-7Rα | Mouse IgG1, κ |
| CD14 | CD14_TotalC | R2 | 5PNNNNNNNNNN(BC) | TCTCAGACCTCCGTA | Antibody Capture | TotalSeq™-C0081 anti-human CD14 Antibody | M5E2 | Monocyte differentiation antigen CD14, myeloid cell-specific leucine-rich glycoprotein, LPS receptor | Mouse IgG2a, κ |
| CD19 | CD19_TotalC | R2 | 5PNNNNNNNNNN(BC) | CTGGGCAATTACTCG | Antibody Capture | TotalSeq™-C0050 anti-human CD19 Antibody | HIB19 | B4 | Mouse IgG1, κ |
| CD197 | CD197_TotalC | R2 | 5PNNNNNNNNNN(BC) | AGTTCAGTCAACCGA | Antibody Capture | TotalSeq™-C0148 anti-human CD197 (CCR7) Antibody | G043H7 | BLR2, CDw197, EBI1, CMKBR7 | Mouse IgG2a, κ |
| CD20 | CD20_TotalC | R2 | 5PNNNNNNNNNN(BC) | TTCTGGGTCCCTAGA | Antibody Capture | TotalSeq™-C0100 anti-human CD20 Antibody | 2H7 | B1, Bp35 | Mouse IgG2b, κ |
| CD279 | CD279_TotalC | R2 | 5PNNNNNNNNNN(BC) | ACAGCGCCGTATTTA | Antibody Capture | TotalSeq™-C0088 anti-human CD279 (PD-1) Antibody | EH12.2H7 | PD-1 | Mouse IgG1, κ |
| CD3 | CD3_TotalC | R2 | 5PNNNNNNNNNN(BC) | CTCATTGTAACTCCT | Antibody Capture | TotalSeq™-C0034 anti-human CD3 Antibody | UCHT1 | T3, CD3ε | Mouse IgG1, κ |
| CD4 | CD4_TotalC | R2 | 5PNNNNNNNNNN(BC) | GAGGTTAGTGATGGA | Antibody Capture | TotalSeq™-C0045 anti-human CD4 Antibody | SK3 | T4, Leu3a | Mouse IgG1, κ |
| CD45 | CD45_TotalC | R2 | 5PNNNNNNNNNN(BC) | TCCCTTGCGATTTAC | Antibody Capture | TotalSeq™-C0048 anti-human CD45 Antibody | 2D1 | Leukocyte Common Antigen (LCA), T200 | Mouse IgG1, κ |
| CD45RA | CD45RA_TotalC | R2 | 5PNNNNNNNNNN(BC) | TCAATCCTTCCGCTT | Antibody Capture | TotalSeq™-C0063 anti-human CD45RA Antibody | HI100 | GP180, L-CA, LCA, LY5, T200, PTPRC | Mouse IgG2b, κ |
| CD45RO | CD45RO_TotalC | R2 | 5PNNNNNNNNNN(BC) | CTCCGAATCATGTTG | Antibody Capture | TotalSeq™-C0087 anti-human CD45RO Antibody | UCHL1 | CD45RO | Mouse IgG2a, κ |
| CD8 | CD8_TotalC | R2 | 5PNNNNNNNNNN(BC) | GCGCAACTTGATGAT | Antibody Capture | TotalSeq™-C0046 anti-human CD8 Antibody | SK1 | T8, Leu2 | Mouse IgG1, κ |
| CD27 | CD27_TotalC | R2 | 5PNNNNNNNNNN(BC) | GCACTCCTGCATGTA | Antibody Capture | TotalSeq™-C0154 anti-human CD27 Antibody | O323 | S152, T14, TNFRSF7 | Mouse IgG1, κ |
| CD49d | CD49d_TotalC | R2 | 5PNNNNNNNNNN(BC) | CCATTCAACTTCCGG | Antibody Capture | TotalSeq™-C0576 anti-human CD49d Antibody | 9F10 | VLA-4 α chain, α4 integrin, Integrin α4 chain, ITGA4 | Mouse IgG1, κ |
| CD62L | CD62L_TotalC | R2 | 5PNNNNNNNNNN(BC) | GTCCCTGCAACTTGA | Antibody Capture | TotalSeq™-C0147 anti-human CD62L Antibody | DREG-56 | L-selectin, LECAM-1, LAM-1, Leu-8, TQ-1 | Mouse IgG1, κ |

**Table S2.** HIV-seq capture sequences

| PrimerID | Capture Sequence | Capture Sequence target |
|---|---|---|
| 10XPreGag | AAGCAGTGGTATCAACGCAGAGTACGGGCGCCACTGCTAGAGA | HIV R-U5-pre-Gag |
| 10XPackaging | AAGCAGTGGTATCAACGCAGAGTACGCACCCATCTCTCTCCTTCTAGC | HIV packaging signal |
| 10XPol | AAGCAGTGGTATCAACGCAGAGTACCAAATTTCTACTAATGCTTTTATTTTTTC | HIV pol |
| 10XRRE | AAGCAGTGGTATCAACGCAGAGTACGTCTGGCCTGTACCGTCAGC | HIV rev response element |
| 10XTat-Rev | AAGCAGTGGTATCAACGCAGAGTACGGATCTGTCTCTGTCTCTCTCTCCACC | HIV tat/rev exon 2 |

**Table S3:** Differentially expressed genes between HIV RNA+ cells vs. HIV RNA- cells during viremia (n=4) determined by gene-wise quasi F-tests on the samples' pseudo-bulked gene counts (adjusted p-value < 0.05 and $\log_2$(fold-change) > 0.25)

| gene | $\log_2$ fold-change | adjusted p-value |
|---|---|---|
| HIV | 15.69572571 | 0 |
| ACTG2 | 3.527094247 | 0.000757928 |
| MPO | 2.902690133 | 6.78E-11 |
| CXCL13 | 2.725862663 | 4.68E-10 |
| RFX8 | 2.709633503 | 0.014524473 |
| AL031056.1 | 2.42193835 | 0.000958085 |
| IGFBP6 | 2.042344964 | 0.043165016 |
| AC007405.3 | 2.005726284 | 0.006614857 |
| AC016074.2 | 1.900844631 | 3.01E-10 |
| FOXD1 | 1.889098251 | 0.006628106 |
| TMEM255A | 1.658360011 | 0.009353443 |
| EAF1-AS1 | 1.599345102 | 0.048492539 |
| CD70 | 1.349785302 | 8.63E-05 |
| CPNE7 | 1.348825797 | 0.034952357 |
| CXCR6 | 1.10906545 | 0.0323038 |
| MTRNR2L8 | 0.924755133 | 4.04E-05 |
| AC006369.1 | 0.832696979 | 0.001089893 |
| SLAMF1 | 0.797841902 | 1.17E-05 |
| CTLA4 | 0.797199828 | 6.18E-06 |
| KLRB1 | 0.735893711 | 1.48E-05 |
| CAMK2N1 | 0.726588581 | 0.036844674 |
| LINC00892 | 0.70168915 | 0.00126818 |
| PLCB1 | 0.665693173 | 0.002425112 |
| MIAT | 0.619307912 | 0.010910194 |
| PTGDR | 0.600866892 | 0.02302089 |
| MAF | 0.552431905 | 0.009533939 |
| SMAD3 | 0.546927258 | 0.011801803 |
| MYBL1 | 0.544902446 | 0.010575692 |
| RAB37 | 0.508532964 | 0.029868192 |
| GPR65 | 0.479850312 | 0.044717322 |
| IKZF3 | 0.47334641 | 0.041022651 |
| GBP5 | 0.471055271 | 0.043677797 |

| | | |
|---|---|---|
| RBM38 | -0.498158524 | 0.048492539 |
| PPP1R15A | -0.512126783 | 0.043677797 |
| RCAN3 | -0.515358858 | 0.036354426 |
| CTSW | -0.522412807 | 0.028900398 |
| SELL | -0.523695944 | 0.025417038 |
| IGLC2 | -0.524237623 | 0.043677797 |
| CD27 | -0.52577507 | 0.028229482 |
| FOS | -0.543792631 | 0.048492539 |
| GZMB | -0.544895934 | 0.043677797 |
| FTL | -0.551199828 | 0.043677797 |
| RGCC | -0.583138439 | 0.028229482 |
| AP3M2 | -0.609603578 | 0.014330057 |
| CLIC3 | -0.613563809 | 0.015487555 |
| ZFP36 | -0.614870293 | 0.005594362 |
| CD7 | -0.630019261 | 0.001512872 |
| CD38 | -0.635451428 | 0.007730269 |
| JUNB | -0.64036523 | 0.007402651 |
| PRKCA | -0.642556322 | 0.034952357 |
| SESN3 | -0.662145548 | 0.021883437 |
| GRASP | -0.699407572 | 0.048562251 |
| RIN3 | -0.700339213 | 0.002241324 |
| IGLC3 | -0.707436323 | 0.013179005 |
| PLCL1 | -0.718055187 | 0.048492539 |
| FGR | -0.725967122 | 0.004904503 |
| MAL | -0.727172251 | 0.000166605 |
| IGHM | -0.754106688 | 0.000413703 |
| HLA-DRB1 | -0.754381068 | 0.000162794 |
| AREG | -0.780445909 | 0.026674366 |
| LSR | -0.79369859 | 0.002919479 |
| FHIT | -0.794145616 | 0.024001436 |
| CEBPD | -0.835132619 | 0.000847956 |
| MCM4 | -0.849960863 | 0.048675448 |
| HLA-DRA | -0.886802389 | 1.12E-05 |
| KLRD1 | -0.90144124 | 8.49E-06 |
| DENND5A | -0.920382989 | 0.033788621 |
| NELL2 | -0.958839882 | 0.000274183 |
| SPINT2 | -0.979233499 | 0.000282892 |
| NCR3 | -0.982088545 | 8.63E-05 |
| TIAM1 | -0.990541892 | 0.00515052 |
| KIR3DL2 | -0.996265714 | 0.022256155 |

| | | |
|---|---|---|
| ARMH1 | -1.008053264 | 3.66E-05 |
| IFITM3 | -1.025943762 | 1.14E-08 |
| KLRK1 | -1.032669156 | 4.36E-07 |
| ACTN1 | -1.051153292 | 0.002810387 |
| HDGFL3 | -1.066011114 | 0.036354426 |
| LEF1 | -1.141468364 | 3.32E-11 |
| IFNGR2 | -1.150685373 | 0.00546889 |
| EDA | -1.156310937 | 0.032009378 |
| LST1 | -1.188126182 | 1.82E-06 |
| LINC00402 | -1.202454951 | 0.025417038 |
| TCEA3 | -1.208674307 | 0.001554463 |
| NCF1 | -1.213390405 | 6.23E-07 |
| LAT2 | -1.242818641 | 0.048560126 |
| YBX3 | -1.274826269 | 0.000648758 |
| LGALS2 | -1.290107259 | 0.041478603 |
| SYK | -1.29822196 | 0.041473816 |
| CFD | -1.315455296 | 0.026559027 |
| SPI1 | -1.31637818 | 1.44E-05 |
| CST3 | -1.324414355 | 3.21E-12 |
| FCGR3A | -1.32917498 | 4.68E-10 |
| SLC40A1 | -1.359726419 | 6.47E-05 |
| ITGAX | -1.398331191 | 0.00267566 |
| SOX4 | -1.418755066 | 0.013405848 |
| DUSP6 | -1.441259118 | 0.012378748 |
| FES | -1.443119712 | 0.041747397 |
| IGLV4-69 | -1.456244654 | 1.24E-11 |
| CD79A | -1.457055856 | 0.00263205 |
| PIK3AP1 | -1.480660426 | 0.000950574 |
| TIMP2 | -1.504090633 | 0.048492539 |
| IKZF2 | -1.543356181 | 1.41E-05 |
| IFI30 | -1.55376862 | 1.03E-17 |
| KLRC1 | -1.562873532 | 0.00296791 |
| NEFL | -1.567991162 | 0.00133106 |
| TRDC | -1.587553053 | 1.27E-12 |
| PECAM1 | -1.59360544 | 9.63E-05 |
| MEF2C | -1.600609822 | 0.008109636 |
| SGK1 | -1.608179229 | 0.003311099 |
| CCR7 | -1.62155278 | 1.27E-15 |
| CXXC5 | -1.640962034 | 0.000845591 |
| NCF2 | -1.647269884 | 0.032310702 |

| | | |
|---|---|---|
| IGLV7-43 | -1.660129467 | 0.010396823 |
| IGFBP7 | -1.674506841 | 0.002174917 |
| IGLV2-8 | -1.678116916 | 1.03E-16 |
| KLRC2 | -1.689090729 | 2.93E-11 |
| NCAM1 | -1.71841561 | 0.022678997 |
| B3GNT7 | -1.725847124 | 0.038585418 |
| TXNDC5 | -1.728436801 | 2.92E-19 |
| AIF1 | -1.739113087 | 8.70E-16 |
| ANKRD55 | -1.779900232 | 0.024409646 |
| CD68 | -1.83651015 | 8.36E-05 |
| HCK | -1.843601865 | 0.022678997 |
| KLRC4 | -1.857414213 | 1.12E-05 |
| KLRC3 | -1.880341144 | 0.000331216 |
| FCER1G | -1.924013037 | 1.44E-20 |
| TMIGD2 | -1.924499214 | 4.82E-05 |
| IGLV7-46 | -1.94905255 | 0.01075927 |
| MNDA | -2.012560764 | 8.00E-06 |
| IGKV1-27 | -2.033249901 | 9.36E-05 |
| PLBD1 | -2.039889333 | 0.031157463 |
| TYROBP | -2.04678287 | 2.34E-29 |
| IGLV1-47 | -2.089494984 | 5.29E-05 |
| TNFSF13 | -2.099452283 | 0.024393201 |
| CLEC7A | -2.105421653 | 0.011149001 |
| IGLV3-19 | -2.148901648 | 9.27E-11 |
| CD300E | -2.164652764 | 0.030043239 |
| IGHV4-59 | -2.189062261 | 4.68E-13 |
| IRF8 | -2.206291624 | 0.001135993 |
| PLXDC2 | -2.210619948 | 0.011670027 |
| FCER1A | -2.211377708 | 0.001117282 |
| IGKV1-12 | -2.218682263 | 0.000332078 |
| CYBB | -2.236037695 | 0.000170704 |
| TRDV1 | -2.250149833 | 9.46E-23 |
| LYZ | -2.298430138 | 1.23E-31 |
| C19orf38 | -2.317179668 | 0.014860317 |
| SH2D1B | -2.338458702 | 0.000163886 |
| IGLV3-9 | -2.429977813 | 3.37E-11 |
| CPVL | -2.43886247 | 6.86E-05 |
| CD14 | -2.458793267 | 0.000105322 |
| MZB1 | -2.458799916 | 1.77E-25 |
| CLDN5 | -2.474246375 | 0.01075927 |

| | | |
|---|---|---|
| AC243829.4 | -2.479402003 | 0.045082413 |
| IGHG2 | -2.489667135 | 1.78E-07 |
| MARCKS | -2.547606444 | 0.000780163 |
| LTBR | -2.561141783 | 0.034952357 |
| MYCN | -2.562768606 | 0.027976369 |
| DAPK1 | -2.572081791 | 0.034952357 |
| TMPPE | -2.630189185 | 0.033788621 |
| KLHL14 | -2.668146313 | 0.020007558 |
| IGKV3-11 | -2.778983796 | 9.05E-45 |
| IGHV4-34 | -2.784813532 | 2.90E-09 |
| AC022075.1 | -2.790249882 | 0.010922577 |
| ZNF385A | -2.81824062 | 0.000491168 |
| TNFAIP2 | -2.858511825 | 0.000282892 |
| ADTRP | -2.860872863 | 6.05E-08 |
| IGKV2-30 | -2.983427595 | 9.16E-14 |
| IGHV3-33 | -3.034015042 | 3.20E-07 |
| IGKV3-15 | -3.047620322 | 4.76E-48 |
| MAFB | -3.05690257 | 0.000277631 |
| CD36 | -3.07581484 | 0.003266843 |
| IGHV3-20 | -3.1665036 | 0.0023283 |
| IGHV1-2 | -3.166714097 | 0.000198388 |
| SIGLEC1 | -3.187200476 | 0.034952357 |
| C1QB | -3.197571291 | 0.001526553 |
| SLC15A3 | -3.216251993 | 0.034047965 |
| IGHV4-4 | -3.243617105 | 0.001296152 |
| SMIM25 | -3.251032339 | 0.034930726 |
| JCHAIN | -3.2682824 | 8.65E-64 |
| LILRA5 | -3.30898375 | 0.000724283 |
| IGLV2-14 | -3.319146564 | 2.29E-68 |
| IGHV1-3 | -3.324470787 | 2.74E-09 |
| IGHV3-64 | -3.329459577 | 9.63E-05 |
| AC233755.1 | -3.363027247 | 0.000513662 |
| IGKV4-1 | -3.377128608 | 1.39E-64 |
| IGKV1-39 | -3.392499812 | 1.21E-19 |
| IGHG1 | -3.392646321 | 5.28E-60 |
| IGHV3-11 | -3.426250174 | 5.19E-06 |
| IGHA2 | -3.43151522 | 4.30E-09 |
| AC233755.2 | -3.467112138 | 4.98E-06 |
| IGKC | -3.481552497 | 4.25E-62 |
| IGLC1 | -3.485070804 | 2.50E-36 |

| | | |
|---|---|---|
| PILRA | -3.519056135 | 0.000168858 |
| MPEG1 | -3.529907226 | 0.000139611 |
| IGHV3-23 | -3.569458297 | 3.07E-66 |
| IGLV6-57 | -3.571529491 | 2.74E-21 |
| SLC7A7 | -3.639759961 | 0.005246075 |
| S100A9 | -3.664920315 | 4.05E-61 |
| IGLV1-44 | -3.669316899 | 3.14E-45 |
| DERL3 | -3.670512305 | 4.55E-09 |
| IGHV3-15 | -3.68837317 | 2.36E-10 |
| IGKV3-20 | -3.695315375 | 3.75E-69 |
| FCGR2A | -3.715433287 | 0.003119319 |
| IGHV5-51 | -3.767083889 | 8.80E-18 |
| IGKV1-5 | -3.781679242 | 3.48E-60 |
| AC020656.1 | -3.785614921 | 0.002297398 |
| CD8B | -3.787133869 | 2.47E-08 |
| CSF1R | -3.791533591 | 0.001867963 |
| IGHV3-53 | -3.796108614 | 8.84E-19 |
| IGKV1-9 | -3.811093444 | 1.06E-34 |
| SERPINA1 | -3.81258581 | 1.40E-13 |
| IGLV8-61 | -3.840289324 | 4.04E-19 |
| IGLV3-25 | -3.855184747 | 2.25E-63 |
| IGLV2-23 | -3.89436899 | 6.21E-73 |
| LILRB2 | -3.911457987 | 4.92E-06 |
| TMEM176A | -3.985084857 | 0.000736078 |
| TNFRSF17 | -4.011320985 | 1.51E-06 |
| VCAN | -4.040812882 | 4.68E-10 |
| S100A12 | -4.062884766 | 3.95E-07 |
| APOBEC3A | -4.07367319 | 4.36E-07 |
| IGLV1-51 | -4.116964922 | 2.25E-63 |
| IGHA1 | -4.141730173 | 3.96E-63 |
| IGLV2-11 | -4.324933696 | 2.83E-69 |
| CSTA | -4.457228104 | 3.82E-05 |
| IGHV3-48 | -4.486676783 | 1.88E-20 |
| IGHV4-39 | -4.572599906 | 2.45E-60 |
| IGKV1-16 | -4.631008989 | 1.14E-21 |
| IGLV3-21 | -4.665723384 | 1.00E-72 |
| IGHV3-7 | -4.692821107 | 1.12E-54 |
| S100A8 | -4.710181242 | 3.39E-72 |
| IGHV1-69D | -4.712802722 | 8.66E-70 |
| IGHV4-31 | -4.73582428 | 2.42E-17 |

| | | |
|---|---|---|
| IGHV1-46 | -4.821050663 | 1.53E-68 |
| IGLV5-37 | -4.843772216 | 1.28E-07 |
| IGKV3D-11 | -4.870045624 | 1.81E-15 |
| IGHV1-69 | -4.908518112 | 5.60E-44 |
| IGHV3-30 | -4.92343077 | 1.92E-85 |
| IGHV4-61 | -4.978065121 | 5.34E-61 |
| FCN1 | -5.198955656 | 4.99E-36 |
| IGKV1-17 | -5.284466828 | 2.26E-43 |
| IGLV3-27 | -5.590746387 | 7.47E-14 |
| IGHV1-18 | -5.6821724 | 1.53E-60 |
| IGLV1-40 | -5.758593743 | 2.85E-104 |
| IGHV5-10-1 | -6.079056189 | 3.63E-15 |
| IGHV3-72 | -6.303147268 | 4.94E-17 |
| PKHD1L1 | -8.724984507 | 0.048492539 |
| RGPD5 | -8.79335847 | 0.038895611 |
| ASGR1 | -8.859005423 | 0.031157463 |
| MYH3 | -8.886900289 | 0.028274397 |
| RNASE2 | -9.032764217 | 0.015358135 |
| KDF1 | -9.054842444 | 0.014043458 |
| FAM157C | -9.141707925 | 0.024878246 |
| CRHBP | -9.202055999 | 0.007180849 |
| GPBAR1 | -9.347762506 | 0.003154004 |
| IGLV4-3 | -9.381980762 | 0.00267566 |
| C1QC | -9.41712207 | 0.002174917 |
| CSF2RA | -9.445382989 | 0.001834245 |
| IGHV3-66 | -9.548723033 | 0.00392874 |
| PLXNB2 | -9.593524603 | 0.000736078 |
| C5AR1 | -9.651288385 | 0.000964068 |
| IGHV1-24 | -10.13600718 | 0.000103782 |
| IGKV2D-28 | -10.28188214 | 5.03E-06 |
| IGHV4-28 | -10.36446117 | 1.24E-05 |
| TMEM176B | -10.46182751 | 9.14E-06 |
| C1QA | -10.61670864 | 1.92E-06 |
| IGHV3-43 | -11.44166474 | 4.61E-10 |
| IGKV1-8 | -12.95759282 | 1.38E-17 |

**Table S4:** Differentially expressed genes between HIV RNA+ cells vs. HIV RNA- cells during ART suppression (n=3) determined by genewise quasi F-tests on the samples' pseudo-bulked gene counts (adjusted p-value < 0.05 and $\log_2$(fold-change) > 0.25)

| gene | $Log_2$ fold-change | adjusted p-value |
|---|---|---|
| HIV | 1.18E-05 | 9.73E-216 |
| SELL | -1.960660595 | 0.010043774 |
| S100A9 | -4.28490229 | 0.032958961 |
| CST3 | -15.29310124 | 0.024718517 |

**Table S5:** Differentially expressed genes in paired samples of CD4+ T cells from during viremia vs. ART suppression (n=3) determined by genewise quasi F-tests on the samples' pseudo-bulked gene counts (p-value < 0.05 and $\log_2$(fold-change) > 0.25)

| gene | $\mathrm{Log_2}$ fold-change | p-value |
|---|---|---|
| IGKV2D-40 | 6.50985737 | 0.00378427 |
| IGKV2-40 | 5.19861084 | 0.00810806 |
| AC099560.1 | 3.54118392 | 0.02689646 |
| IGHV3-64D | 3.29686926 | 0.00521642 |
| CLDN1 | 3.16933682 | 0.0148928 |
| LINC02206 | 3.12404319 | 0.01134247 |
| FIBCD1 | 3.02397365 | 0.01003443 |
| AL109910.2 | 2.92597255 | 0.0055532 |
| PRTN3 | 2.8273881 | 0.01570934 |
| DPP6 | 2.55791016 | 0.01884257 |
| NCKAP5 | 2.45656729 | 0.01511861 |
| PSMB11 | 2.40193626 | 0.0414188 |
| ZNF30-AS1 | 2.39161906 | 0.02003295 |
| DSC1 | 2.33209652 | 0.00784307 |
| DBH | 2.30485428 | 0.03927556 |
| PPY | 2.30013978 | 0.01431054 |
| GDF10 | 2.27252195 | 0.00193903 |
| XCR1 | 2.21628364 | 0.01085843 |
| AC093390.2 | 2.21479521 | 0.03860464 |
| PCAT19 | 2.1467278 | 0.01858777 |
| AL161644.1 | 2.09299043 | 0.01267603 |
| AC114291.2 | 2.07206686 | 0.02584894 |
| FP236315.1 | 2.05074231 | 0.01425985 |
| DACT1 | 2.02675206 | 0.00851737 |
| AC005580.1 | 1.96401487 | 0.01610625 |
| BX322557.2 | 1.96369582 | 0.0052297 |
| AC108136.1 | 1.95170315 | 0.03130457 |
| RASL10B | 1.9449411 | 0.01070363 |
| LCN10 | 1.92117093 | 0.01065068 |
| DNASE1L3 | 1.9060923 | 0.00675722 |
| LINC01281 | 1.88610321 | 0.00595718 |
| NBAT1 | 1.88043298 | 0.02489305 |

| | | |
|---|---|---|
| DDIT4L | 1.87964732 | 0.01247171 |
| AJ011931.2 | 1.87570236 | 0.04066183 |
| ZNF233 | 1.86287616 | 0.02330544 |
| AJAP1 | 1.84371316 | 0.03502714 |
| XKRX | 1.82099675 | 0.0419237 |
| SLC28A2 | 1.7843616 | 0.02371159 |
| LINC02295 | 1.73536717 | 0.0184731 |
| AC093418.1 | 1.71939586 | 0.02444076 |
| AL158151.1 | 1.70913165 | 0.01674694 |
| CA6 | 1.68389913 | 0.02359607 |
| EDAR | 1.65933724 | 0.01940173 |
| AC078883.2 | 1.6470167 | 0.03107244 |
| AC233976.2 | 1.63417656 | 0.01591677 |
| AC087354.1 | 1.63313986 | 0.04456227 |
| AC103843.1 | 1.62465896 | 0.01609594 |
| SORCS3 | 1.616489 | 0.02253232 |
| ADD2 | 1.60856341 | 0.00492278 |
| AP001011.1 | 1.60575641 | 0.0384398 |
| AC053503.2 | 1.59853405 | 0.04197931 |
| AC129507.1 | 1.54884561 | 0.04506098 |
| VMO1 | 1.52527637 | 0.03515923 |
| AL357153.1 | 1.51518305 | 0.04751264 |
| LINC01624 | 1.50788893 | 0.03892362 |
| AP004609.1 | 1.50256882 | 0.0159108 |
| TMEM272 | 1.49769415 | 0.02692149 |
| AC008892.1 | 1.4498805 | 0.04468274 |
| ZNF285 | 1.44894085 | 0.04263204 |
| ZSCAN23 | 1.44623264 | 0.02655838 |
| CELA1 | 1.44426582 | 0.01595209 |
| MMP28 | 1.44115179 | 0.04267445 |
| AC139795.3 | 1.42210387 | 0.03576317 |
| SH3PXD2B | 1.41798551 | 0.02395946 |
| GTSCR1 | 1.38938055 | 0.02980386 |
| ZNF157 | 1.38557328 | 0.01394491 |
| AC006372.1 | 1.37774998 | 0.03712831 |
| RAB7B | 1.37575593 | 0.03958968 |
| CR559946.1 | 1.37500863 | 0.03200932 |
| AL118511.4 | 1.36425145 | 0.03200046 |
| AC018462.1 | 1.34627229 | 0.02992338 |
| CR2 | 1.34596769 | 0.03504037 |

| | | |
|---|---|---|
| MCF2L-AS1 | 1.34109274 | 0.02477448 |
| MEOX1 | 1.33562427 | 0.01595627 |
| SULT1B1 | 1.33487353 | 0.00522374 |
| NODAL | 1.33312274 | 0.02038736 |
| GAL3ST4 | 1.32656549 | 0.02111714 |
| AC006504.1 | 1.28959696 | 0.04866319 |
| AC083870.1 | 1.28765666 | 0.04568935 |
| GPC3 | 1.28635328 | 0.01481361 |
| KLHL34 | 1.26962519 | 0.00980339 |
| SMARCA1 | 1.26753869 | 0.01492121 |
| GPA33 | 1.26719749 | 0.0120959 |
| ANKRD55 | 1.26313213 | 0.01329197 |
| AC010746.2 | 1.24383739 | 0.02343516 |
| SLC5A10 | 1.24339481 | 0.01101092 |
| LINC01709 | 1.2413964 | 0.04743458 |
| MMEL1 | 1.23886231 | 0.02360198 |
| DBNDD1 | 1.23244568 | 0.00953 |
| MYO1B | 1.22223277 | 0.02268973 |
| SEMA3A | 1.21822461 | 0.03236168 |
| ZNF32-AS1 | 1.20509232 | 0.02370612 |
| AC018797.3 | 1.20384013 | 0.04487525 |
| AC245884.8 | 1.20010405 | 0.0350388 |
| AC093535.1 | 1.19694725 | 0.02778518 |
| AEBP1 | 1.19672579 | 0.02403423 |
| SPATA32 | 1.19382837 | 0.03197167 |
| GRAPL | 1.18059645 | 0.02479329 |
| AL590822.2 | 1.17322981 | 0.03750325 |
| AC117503.5 | 1.16087039 | 0.04417971 |
| AL139106.1 | 1.15952363 | 0.04325742 |
| AC091965.1 | 1.15140008 | 0.04364821 |
| EPHA1 | 1.14481527 | 0.03179036 |
| CNNM1 | 1.142865 | 0.04289862 |
| AP002755.1 | 1.1414268 | 0.02824682 |
| PRKCA-AS1 | 1.13770582 | 0.02277117 |
| KCNH2 | 1.1237758 | 0.00728843 |
| EVPL | 1.09609419 | 0.03963551 |
| ASB15 | 1.08947596 | 0.04563223 |
| ADTRP | 1.08564828 | 0.02210854 |
| BCDIN3D-AS1 | 1.0727484 | 0.02731549 |
| AC140125.2 | 1.06988841 | 0.04649444 |

| | | |
|---|---|---|
| AS3MT | 1.06908263 | 0.0336049 |
| SARDH | 1.06836451 | 0.00562428 |
| AC011491.3 | 1.06373861 | 0.00915805 |
| GAS5-AS1 | 1.05935079 | 0.01689292 |
| DNAH6 | 1.05894797 | 0.02327219 |
| AC103563.9 | 1.05485869 | 0.03650795 |
| LINC01727 | 1.05015527 | 0.048415 |
| PNMA8B | 1.04920912 | 0.04849151 |
| NPAS2 | 1.04893369 | 0.04922576 |
| AC008462.1 | 1.04837454 | 0.04965057 |
| NUPR2 | 1.04776582 | 0.02317093 |
| AL035252.2 | 1.04248016 | 0.04920149 |
| FGFR2 | 1.04170703 | 0.0338871 |
| ACTN1 | 1.03831399 | 0.01162011 |
| AC015921.1 | 1.03371747 | 0.04154774 |
| AC008752.2 | 1.02892387 | 0.03746357 |
| GSDMA | 1.02594326 | 0.01884121 |
| GRTP1 | 1.02555979 | 0.04714412 |
| MARVELD2 | 1.02545051 | 0.03957834 |
| AL034428.1 | 1.02313368 | 0.0149394 |
| CHGB | 1.02078359 | 0.04559358 |
| LINC01063 | 1.01588944 | 0.0175219 |
| LINC01891 | 1.006152 | 0.00900559 |
| TXNRD3 | 1.00259184 | 0.01700661 |
| LINC00402 | 0.99534357 | 0.01398405 |
| LEF1-AS1 | 0.99482502 | 0.02373047 |
| NCMAP-DT | 0.99324385 | 0.00395203 |
| AC005842.1 | 0.98748616 | 0.02619375 |
| NUDT12 | 0.9793338 | 0.02752876 |
| AL117381.1 | 0.97732607 | 0.04367988 |
| RNF157-AS1 | 0.97645962 | 0.02118132 |
| DTX1 | 0.97112123 | 0.00905717 |
| AL133383.1 | 0.96716089 | 0.0485868 |
| AC073834.1 | 0.95972192 | 0.0293221 |
| TEAD2 | 0.95958106 | 0.04097446 |
| AL138963.4 | 0.95864828 | 0.0052103 |
| MDS2 | 0.95852138 | 0.02204306 |
| ZNF662 | 0.95338822 | 0.01915196 |
| AC124068.2 | 0.95100913 | 0.02326546 |
| ARG1 | 0.94129236 | 0.0427278 |

| | | |
|---|---|---|
| AC113143.3 | 0.93455357 | 0.03448981 |
| ATG9B | 0.93177671 | 0.00941305 |
| LINC01725 | 0.9313883 | 0.04725208 |
| EPPK1 | 0.9286018 | 0.02347561 |
| AC008443.5 | 0.92420596 | 0.01120683 |
| GRAP | 0.91887524 | 0.03891959 |
| AC138028.2 | 0.91425084 | 0.03834727 |
| IL6R | 0.89322761 | 0.03813332 |
| AL353660.1 | 0.88619948 | 0.0151523 |
| AC108488.1 | 0.885321 | 0.04305644 |
| ZNF793-AS1 | 0.88466194 | 0.01898389 |
| LIMS2 | 0.88313281 | 0.02274685 |
| PRKAG2-AS1 | 0.88143341 | 0.03295399 |
| KALRN | 0.87736768 | 0.03954301 |
| ASB13 | 0.8752236 | 0.04132471 |
| AC010896.1 | 0.87392351 | 0.04957844 |
| ACER1 | 0.87382799 | 0.04094574 |
| TMIGD2 | 0.87161949 | 0.01612867 |
| TNFRSF10D | 0.8713898 | 0.0189425 |
| CHRM3-AS2 | 0.86939676 | 0.02628265 |
| AC109347.1 | 0.86694608 | 0.04842882 |
| LINC01259 | 0.8659502 | 0.04598015 |
| LINC00866 | 0.86282375 | 0.01489174 |
| CCR7 | 0.8615396 | 0.02634098 |
| AJUBA | 0.85209545 | 0.04900668 |
| AC003086.1 | 0.84586983 | 0.04524844 |
| LINC02714 | 0.84556103 | 0.04873663 |
| RETREG1 | 0.84545824 | 0.01360898 |
| ZNF582-AS1 | 0.84084969 | 0.01546057 |
| OSER1-DT | 0.83456623 | 0.01825182 |
| S100B | 0.82600923 | 0.03256153 |
| EPHX2 | 0.823775 | 0.02552387 |
| LDLRAP1 | 0.82264664 | 0.03332369 |
| AP000977.1 | 0.82170953 | 0.0354419 |
| VSIG1 | 0.81629422 | 0.02042913 |
| AC015712.1 | 0.81363113 | 0.0479665 |
| ZNF667-AS1 | 0.80817799 | 0.02465758 |
| AC022706.1 | 0.80238655 | 0.04817511 |
| AK5 | 0.80111476 | 0.03592291 |
| AL163973.2 | 0.80097886 | 0.04693652 |

| | | |
|---|---|---|
| LEF1 | 0.79983762 | 0.02275728 |
| KRT73 | 0.79683789 | 0.04694071 |
| NEXMIF | 0.79598324 | 0.02232049 |
| SLC12A9-AS1 | 0.79282346 | 0.02344535 |
| DNAJC27-AS1 | 0.77964625 | 0.01511056 |
| CDK20 | 0.77311567 | 0.03911118 |
| AC007342.4 | 0.77262879 | 0.03764368 |
| LINC01550 | 0.77081497 | 0.01907887 |
| EPHA1-AS1 | 0.76749784 | 0.01219062 |
| TMEM220 | 0.76645333 | 0.0211762 |
| GCSAM | 0.76633191 | 0.02675904 |
| BX293535.1 | 0.76308214 | 0.01383644 |
| ZNF284 | 0.75773959 | 0.04777111 |
| AC106707.1 | 0.75770967 | 0.04745475 |
| AL158055.1 | 0.75599438 | 0.04500678 |
| AL592211.2 | 0.75310425 | 0.04933 |
| RPS10 | 0.75020249 | 0.00476845 |
| PCBP3 | 0.7464726 | 0.02823881 |
| TTC9 | 0.74480565 | 0.01610295 |
| LINC02762 | 0.74259252 | 0.01528529 |
| AC103563.7 | 0.73587382 | 0.02394289 |
| MTUS1 | 0.72605019 | 0.0256079 |
| GCNT4 | 0.72402534 | 0.01168339 |
| DCBLD2 | 0.72320661 | 0.04723059 |
| AC253572.2 | 0.72230424 | 0.01157856 |
| ITGA6 | 0.71359591 | 0.01674925 |
| GNAI1 | 0.71215324 | 0.0326997 |
| ALS2CL | 0.70612466 | 0.04409488 |
| AC013264.1 | 0.7019285 | 0.04016556 |
| C16orf74 | 0.70099077 | 0.02871197 |
| AC011450.1 | 0.69416306 | 0.03039114 |
| SAXO2 | 0.6936115 | 0.0235607 |
| SLC1A7 | 0.69036221 | 0.02511314 |
| TMC3-AS1 | 0.68689523 | 0.04424258 |
| VIPR1 | 0.68357339 | 0.02157544 |
| Z93241.1 | 0.67721347 | 0.01233805 |
| CRIP2 | 0.67510635 | 0.02294125 |
| AC139720.1 | 0.67279314 | 0.04291961 |
| CCDC30 | 0.67089142 | 0.04149394 |
| AC025171.2 | 0.67006246 | 0.03330041 |

| | | |
|---|---|---|
| AL390198.1 | 0.66991519 | 0.04268014 |
| AL136038.5 | 0.66930486 | 0.01829464 |
| ISM1 | 0.66884904 | 0.01751164 |
| AC012146.1 | 0.66520318 | 0.01488985 |
| RCN3 | 0.65865413 | 0.03405516 |
| TCF7 | 0.65615602 | 0.02658557 |
| ARHGEF10 | 0.65442277 | 0.04381649 |
| FAM153CP | 0.6468694 | 0.01603143 |
| PAXBP1-AS1 | 0.63504588 | 0.03849294 |
| PLEKHB1 | 0.63078218 | 0.02877007 |
| AMPD3 | 0.62860219 | 0.04631597 |
| AKR1E2 | 0.62511876 | 0.0487274 |
| AC020922.4 | 0.62390684 | 0.045545 |
| FBLN7 | 0.61494044 | 0.01597625 |
| RCAN3AS | 0.61449965 | 0.04737621 |
| SH2B1 | 0.60966654 | 0.02604879 |
| SNPH | 0.60241559 | 0.02906445 |
| TCEA3 | 0.6019178 | 0.02457169 |
| ZNF154 | 0.58325744 | 0.04516395 |
| CRTC1 | 0.5816472 | 0.03303927 |
| NEK11 | 0.57836525 | 0.03522554 |
| STMN3 | 0.57633123 | 0.01302868 |
| PKIA | 0.57580517 | 0.01800087 |
| RFLNB | 0.57461737 | 0.02431645 |
| AL021155.5 | 0.57059176 | 0.0395856 |
| AC007952.4 | 0.56951729 | 0.01363359 |
| GOLGA7B | 0.56821934 | 0.01825558 |
| MAL | 0.56602765 | 0.01502806 |
| RGL4 | 0.56479311 | 0.03764002 |
| ZNF439 | 0.56463427 | 0.02694334 |
| INPP4B | 0.56343086 | 0.03687337 |
| EPHB6 | 0.55895764 | 0.02806264 |
| AC245014.3 | 0.55732761 | 0.02814137 |
| LINC01011 | 0.55655399 | 0.03791158 |
| PHOSPHO2 | 0.55182523 | 0.04569584 |
| TMEM191B | 0.5460799 | 0.0218435 |
| LINC02361 | 0.54151067 | 0.04942103 |
| CSGALNACT1 | 0.53662753 | 0.03574837 |
| MEGF6 | 0.53481249 | 0.03673686 |
| MT-ND5 | 0.53283569 | 0.04212225 |

| | | |
|---|---|---|
| AL451085.1 | 0.53218769 | 0.03835649 |
| TPCN1 | 0.5319924 | 0.04470961 |
| C10orf143 | 0.51692899 | 0.04772706 |
| ZBED5 | 0.51667574 | 0.03517244 |
| SUSD4 | 0.51468233 | 0.03161692 |
| AC008555.4 | 0.51373078 | 0.02484258 |
| NOXA1 | 0.51278169 | 0.04562114 |
| TBC1D17 | 0.51185348 | 0.04752635 |
| SLC39A4 | 0.51075786 | 0.03860683 |
| SNRPN | 0.50716691 | 0.03595567 |
| SAMD10 | 0.50442624 | 0.03159381 |
| MT-CO1 | 0.50401438 | 0.04552494 |
| MFNG | 0.50014847 | 0.03476208 |
| PPP1R3F | 0.49966086 | 0.03117253 |
| NOSIP | 0.49780256 | 0.04815789 |
| MT-ND4 | 0.49566308 | 0.03528252 |
| OXNAD1 | 0.49527547 | 0.04248531 |
| ST6GALNAC1 | 0.49356028 | 0.04977354 |
| SNHG10 | 0.48711408 | 0.04458458 |
| MAP3K12 | 0.48015929 | 0.04926393 |
| AC073896.2 | 0.47154697 | 0.04734492 |
| HSPB1 | 0.47102613 | 0.02302157 |
| FAM117A | 0.46404502 | 0.04742441 |
| ITGB2-AS1 | 0.46032048 | 0.02788313 |
| AC010642.2 | 0.45476823 | 0.03870929 |
| ZNF33B | 0.45475531 | 0.04340844 |
| CD7 | 0.44754234 | 0.04399284 |
| MAP11 | 0.44556655 | 0.04618547 |
| AC027644.3 | 0.44535473 | 0.04089112 |
| PRMT2 | 0.44043004 | 0.03644567 |
| PRKCQ-AS1 | 0.43938028 | 0.03849294 |
| ATP6V0E2 | 0.41703944 | 0.03672977 |
| SPINT2 | 0.40733058 | 0.04538944 |
| PSMA2 | -0.3829118 | 0.04847947 |
| HSPE1 | -0.3853497 | 0.04755836 |
| MRPL17 | -0.395032 | 0.04981018 |
| TUBA1C | -0.4090422 | 0.04807691 |
| NEAT1 | -0.4103997 | 0.04122022 |
| PLPP5 | -0.4135052 | 0.04647181 |
| TSG101 | -0.425271 | 0.04664641 |

| | | |
|---|---|---|
| USP1 | -0.4273196 | 0.03433342 |
| HMGA1 | -0.4285928 | 0.04683511 |
| SLBP | -0.4288055 | 0.03801571 |
| SARNP | -0.4353576 | 0.0471051 |
| YWHAE | -0.4441481 | 0.03964697 |
| NSD2 | -0.4466028 | 0.04584062 |
| TOP1 | -0.461382 | 0.03668881 |
| WDFY1 | -0.468338 | 0.0291949 |
| SP100 | -0.4778156 | 0.03962389 |
| SLAMF1 | -0.4788235 | 0.02887687 |
| CACYBP | -0.4827066 | 0.0332202 |
| MRPL13 | -0.482996 | 0.02489595 |
| LRRC42 | -0.4860459 | 0.04183327 |
| IL2RB | -0.4888798 | 0.03305418 |
| IFIT5 | -0.4931448 | 0.02266916 |
| PGRMC1 | -0.4977421 | 0.03322184 |
| HCCS | -0.4979679 | 0.0402272 |
| ALYREF | -0.4980881 | 0.03774971 |
| CHPF | -0.49858 | 0.03400391 |
| PDXK | -0.4994091 | 0.02328617 |
| CDK2AP1 | -0.5023671 | 0.04561489 |
| POMP | -0.5079539 | 0.03764823 |
| MFAP1 | -0.5106706 | 0.04659632 |
| RPA3 | -0.5112266 | 0.04304804 |
| SNRPG | -0.5139767 | 0.02095795 |
| TMEM140 | -0.5141332 | 0.03958311 |
| TM2D2 | -0.5184419 | 0.0357417 |
| MRPL22 | -0.5196921 | 0.04802629 |
| INPP1 | -0.5200814 | 0.03804259 |
| MICB | -0.5211463 | 0.03117555 |
| NDUFV2 | -0.5309783 | 0.04936283 |
| EIF4A3 | -0.5313705 | 0.03091824 |
| GZMB | -0.5369625 | 0.02673896 |
| MAD2L1 | -0.5471592 | 0.02537888 |
| BAK1 | -0.5509399 | 0.04785676 |
| CARHSP1 | -0.5519311 | 0.02518187 |
| DSN1 | -0.555934 | 0.03428594 |
| TUBB | -0.5604543 | 0.04528715 |
| NAA50 | -0.5606305 | 0.02606623 |
| CBLB | -0.5640335 | 0.0478281 |

| | | |
|---|---|---|
| PGAP1 | -0.5676075 | 0.02249435 |
| SH2D1A | -0.5704401 | 0.02134408 |
| DONSON | -0.5735102 | 0.02655729 |
| CDC7 | -0.5751323 | 0.02961059 |
| CLDND1 | -0.5841997 | 0.04275285 |
| HIST1H2AC | -0.5897106 | 0.02628971 |
| PSMA3 | -0.5898746 | 0.01171858 |
| TUBB4B | -0.593189 | 0.02051375 |
| SLC35B1 | -0.5935147 | 0.02224606 |
| GINS4 | -0.5958465 | 0.02764681 |
| TMEM170B | -0.5962995 | 0.04161379 |
| BARD1 | -0.6039296 | 0.02608802 |
| TLR2 | -0.6047313 | 0.04024473 |
| HIST1H4C | -0.6048832 | 0.00998085 |
| RAB33A | -0.6098278 | 0.04770468 |
| TXNDC17 | -0.6104532 | 0.01320299 |
| N4BP1 | -0.6113226 | 0.04622995 |
| HAUS8 | -0.6176777 | 0.03665494 |
| IFNLR1 | -0.6194092 | 0.03923707 |
| PHF19 | -0.6246335 | 0.01490252 |
| CALCB | -0.6266001 | 0.04984432 |
| H2AFZ | -0.6273822 | 0.0160387 |
| HIST1H2AG | -0.6314783 | 0.02592281 |
| SMC4 | -0.6338477 | 0.01670921 |
| HIST1H4D | -0.6411825 | 0.03870905 |
| AC147651.4 | -0.6418403 | 0.04019261 |
| HELLS | -0.6430296 | 0.01463404 |
| ZNF367 | -0.6432692 | 0.04385962 |
| TCF7L2 | -0.6435492 | 0.04167887 |
| LRR1 | -0.6514096 | 0.01017229 |
| HIST2H2BF | -0.6536482 | 0.0234456 |
| PARPBP | -0.6539329 | 0.0197282 |
| CDCA4 | -0.6552528 | 0.0150814 |
| TPM4 | -0.6560504 | 0.01374679 |
| YBX3 | -0.6563402 | 0.04963363 |
| FANCL | -0.6569114 | 0.03854934 |
| MYO16 | -0.664653 | 0.04782439 |
| ECHDC3 | -0.6694654 | 0.04610008 |
| FAM83D | -0.6728736 | 0.02563716 |
| PTMS | -0.6734264 | 0.00665035 |

| | | |
|---|---|---|
| HLA-DMB | -0.6787183 | 0.04107029 |
| SDC4 | -0.6802315 | 0.04639472 |
| WARS | -0.6833833 | 0.01044351 |
| HIST1H2BI | -0.6847286 | 0.04160748 |
| TIMP2 | -0.6851505 | 0.02445577 |
| HIST1H2BB | -0.6901505 | 0.02589027 |
| ATAD5 | -0.6918439 | 0.01099355 |
| HIST1H4H | -0.6919158 | 0.02924069 |
| PIK3AP1 | -0.6936076 | 0.02956544 |
| HES4 | -0.7020729 | 0.01605648 |
| GOLM1 | -0.7062599 | 0.02409392 |
| YWHAH | -0.7122063 | 0.01263115 |
| MIR4435-2HG | -0.7143914 | 0.01201833 |
| ZBP1 | -0.7143986 | 0.00457513 |
| NETO2 | -0.715616 | 0.03153703 |
| MTHFD2 | -0.7225355 | 0.00852814 |
| CLECL1 | -0.728665 | 0.03028166 |
| FKBPL | -0.7304859 | 0.03977634 |
| ARG2 | -0.7315092 | 0.04075115 |
| F2R | -0.7315304 | 0.01608841 |
| LY6E | -0.7330041 | 0.02017824 |
| TUBA1B | -0.7355435 | 0.00897111 |
| RGS16 | -0.7396027 | 0.04957905 |
| FAM27C | -0.7456835 | 0.02013644 |
| AC011379.2 | -0.748277 | 0.03826544 |
| LYST | -0.7528358 | 0.01368309 |
| TNK2-AS1 | -0.7546731 | 0.02304561 |
| SMAD1 | -0.7584621 | 0.04118516 |
| CCDC50 | -0.7612045 | 0.00921016 |
| GRAMD1C | -0.7666206 | 0.04857307 |
| CRACR2B | -0.7696081 | 0.03962167 |
| TP53I3 | -0.7703564 | 0.02418144 |
| PRELID3A | -0.7716936 | 0.04764955 |
| TICAM2 | -0.7729353 | 0.03573205 |
| LGALS9C | -0.7739686 | 0.0398197 |
| RRM1 | -0.7764976 | 0.00480082 |
| FCGR2B | -0.7770691 | 0.01428172 |
| LINC00937 | -0.778378 | 0.03831571 |
| AKAP5 | -0.782801 | 0.04643579 |
| IFIH1 | -0.7836418 | 0.00971376 |

| | | |
|---|---|---|
| PPIF | -0.7869443 | 0.03107169 |
| LINC02413 | -0.7871666 | 0.04394559 |
| KLF4 | -0.7881754 | 0.03798673 |
| SECTM1 | -0.7930052 | 0.02084428 |
| CXXC5 | -0.7936973 | 0.01055789 |
| HIST2H2BE | -0.7953632 | 0.04218035 |
| HIST1H2AI | -0.795684 | 0.01590574 |
| CCNE1 | -0.7996323 | 0.01094998 |
| CMTM1 | -0.8026238 | 0.03706058 |
| PLCG2 | -0.8060215 | 0.04822972 |
| ATAD2 | -0.806035 | 0.03976786 |
| AC034238.1 | -0.8107599 | 0.03895936 |
| MYL6B | -0.8108782 | 0.00796409 |
| LYN | -0.812475 | 0.02992935 |
| IRF4 | -0.8140054 | 0.00683029 |
| BRIP1 | -0.8146004 | 0.01926136 |
| UBE2S | -0.8157751 | 0.01721234 |
| RASGRP3 | -0.8166862 | 0.0142295 |
| CHAF1A | -0.8372177 | 0.00488426 |
| HIST1H3J | -0.838929 | 0.02984066 |
| SLC31A1 | -0.8390148 | 0.04527905 |
| SPHK1 | -0.8393339 | 0.02546196 |
| HIST1H3H | -0.8415282 | 0.00350533 |
| NDC80 | -0.8440131 | 0.00479781 |
| WDR11-AS1 | -0.8444217 | 0.03610621 |
| SLC43A2 | -0.8476724 | 0.04393095 |
| IFI27L1 | -0.8501656 | 0.00786121 |
| ARHGAP11A | -0.8528512 | 0.00613015 |
| DHFR | -0.8535669 | 0.02756233 |
| SLC1A4 | -0.8557993 | 0.01258675 |
| MCM5 | -0.8576921 | 0.01457379 |
| AL445228.2 | -0.8687977 | 0.04535205 |
| TLCD1 | -0.8719809 | 0.03584395 |
| HIST1H2BL | -0.8730304 | 0.02703123 |
| PROB1 | -0.874557 | 0.0401667 |
| PRTFDC1 | -0.8756886 | 0.02298649 |
| CENPI | -0.877057 | 0.04812047 |
| IL12RB2 | -0.8772323 | 0.01890074 |
| PRR11 | -0.8819801 | 0.0110062 |
| SLC31A2 | -0.8827718 | 0.0179116 |

| | | |
|---|---|---|
| MCM7 | -0.8832738 | 0.0108173 |
| MXD3 | -0.8893559 | 0.0062574 |
| HMGB2 | -0.8921769 | 0.00573811 |
| MFSD2A | -0.8923699 | 0.02139711 |
| CENPP | -0.8929337 | 0.00588082 |
| MOB3B | -0.897621 | 0.04992601 |
| LGALS9 | -0.8988765 | 0.02354065 |
| NAPSA | -0.8994006 | 0.03463767 |
| ENTPD1 | -0.9017317 | 0.02283109 |
| YPEL4 | -0.906325 | 0.03531382 |
| GZMK | -0.9079219 | 0.00497704 |
| DOCK4 | -0.91066 | 0.04519832 |
| TYROBP | -0.9120628 | 0.03236662 |
| BRCA1 | -0.9152657 | 0.0409961 |
| CIP2A | -0.9163694 | 0.01047473 |
| ADAMTSL4 | -0.9198579 | 0.02858846 |
| AC104078.1 | -0.9205842 | 0.04371706 |
| H2AFX | -0.921544 | 0.00429124 |
| KIF18A | -0.9243365 | 0.04081605 |
| TCN2 | -0.9256114 | 0.01909203 |
| ABTB2 | -0.9296808 | 0.03839255 |
| LINC01023 | -0.9332809 | 0.03662487 |
| E2F2 | -0.9360111 | 0.01924671 |
| TREX1 | -0.9361518 | 0.0184158 |
| NLRP3 | -0.9364524 | 0.01466074 |
| RHOB | -0.9377059 | 0.02341287 |
| HBEGF | -0.9394101 | 0.03353688 |
| FANCI | -0.9402881 | 0.00658642 |
| PRAM1 | -0.9430378 | 0.02386279 |
| GFOD1 | -0.9493809 | 0.00875009 |
| FAM49A | -0.9495777 | 0.00771274 |
| MS4A1 | -0.9504445 | 0.01101114 |
| CHAF1B | -0.9517574 | 0.00955558 |
| BCL11A | -0.9549907 | 0.0210344 |
| U62317.1 | -0.9624785 | 0.01586747 |
| LRG1 | -0.9650266 | 0.02927991 |
| CSF1R | -0.9666107 | 0.02926946 |
| FEN1 | -0.9687823 | 0.00196081 |
| XRCC2 | -0.9718252 | 0.04287749 |
| LAP3 | -0.9802967 | 0.00691446 |

| | | |
|---|---|---|
| PCNA | -0.9944602 | 0.02036459 |
| GPT2 | -0.9977136 | 0.0352866 |
| CCR5AS | -0.9978568 | 0.02551882 |
| PLK4 | -0.9982279 | 0.00892803 |
| TUBB3 | -1.0010521 | 0.01626675 |
| TOX2 | -1.0036004 | 0.03135035 |
| CDKN2C | -1.0071616 | 0.00553201 |
| SLC41A2 | -1.0080733 | 0.03527307 |
| STAC3 | -1.0091474 | 0.01300784 |
| ABCA1 | -1.0106532 | 0.02729678 |
| AC092718.4 | -1.0136783 | 0.03498688 |
| AC090559.1 | -1.0159805 | 0.0473283 |
| EIF2AK2 | -1.0181452 | 0.01187337 |
| KCNK7 | -1.0195483 | 0.02531318 |
| PLPP2 | -1.0214777 | 0.01341755 |
| DUSP6 | -1.0216079 | 0.01503513 |
| LMNB1 | -1.0246475 | 0.00279877 |
| LPCAT2 | -1.0253997 | 0.03604563 |
| DDR2 | -1.0259242 | 0.02886091 |
| SLC7A7 | -1.0300151 | 0.04570067 |
| AC017002.1 | -1.0303553 | 0.03072853 |
| HIST2H2AA4 | -1.0304131 | 0.04865312 |
| AC144831.1 | -1.0328766 | 0.04728543 |
| HIST1H2AM | -1.036177 | 0.00996345 |
| KPNA2 | -1.0362373 | 0.0364653 |
| PHLDA2 | -1.0395495 | 0.01670456 |
| EZH2 | -1.039946 | 0.01694919 |
| SCO2 | -1.0492991 | 0.01429446 |
| HIST1H2BF | -1.0497934 | 0.00476453 |
| DDIAS | -1.0537602 | 0.00962032 |
| POGLUT2 | -1.054586 | 0.00904093 |
| RECQL4 | -1.0581688 | 0.01280431 |
| MT1E | -1.0593447 | 0.01708015 |
| FCRLA | -1.0597551 | 0.03862267 |
| OIP5 | -1.0609651 | 0.03969384 |
| REXO5 | -1.0630143 | 0.0084086 |
| AC009522.1 | -1.0645469 | 0.03664248 |
| CD70 | -1.066053 | 0.0072064 |
| LINC01943 | -1.0676891 | 0.00660241 |
| AATK | -1.0706185 | 0.0284223 |

| | | |
|---|---|---|
| FBXO5 | -1.0735435 | 0.00193352 |
| SLC1A2 | -1.0748395 | 0.03070013 |
| GGH | -1.0754044 | 0.0040852 |
| MT2A | -1.0759381 | 0.00336062 |
| MCM4 | -1.0762634 | 0.00494251 |
| CTXN1 | -1.0831799 | 0.04063138 |
| HSH2D | -1.0846077 | 0.00230337 |
| MCM6 | -1.0888184 | 0.00286725 |
| CD300LF | -1.0900169 | 0.02176235 |
| ARNTL2 | -1.0919427 | 0.03551967 |
| ALDH1L2 | -1.1000466 | 0.04593727 |
| CORO1C | -1.1023476 | 0.00167772 |
| FCER2 | -1.1041138 | 0.02802206 |
| SLC37A2 | -1.1068369 | 0.02525645 |
| LILRB2 | -1.1070514 | 0.04676796 |
| STMN1 | -1.109481 | 0.00353102 |
| NT5DC2 | -1.1142707 | 0.01181765 |
| SYK | -1.1154821 | 0.01595363 |
| ARHGAP42 | -1.115916 | 0.02683559 |
| GAPT | -1.1191361 | 0.0171345 |
| BCAT1 | -1.1200474 | 0.01079256 |
| TCF4 | -1.1200804 | 0.00749248 |
| AC007569.1 | -1.1275489 | 0.00898082 |
| HIST1H2BJ | -1.1287866 | 0.00275503 |
| LINC02249 | -1.1307246 | 0.01270292 |
| AL592146.2 | -1.1384717 | 0.03737295 |
| AP005899.1 | -1.14029 | 0.0307761 |
| TP63 | -1.1411759 | 0.02735874 |
| AC073878.1 | -1.1433188 | 0.03365483 |
| NDST1 | -1.1444791 | 0.01496633 |
| MUCL1 | -1.1449185 | 0.04144829 |
| SPATS2L | -1.1464199 | 0.00709139 |
| NEIL3 | -1.1466552 | 0.0263519 |
| CD79A | -1.1523405 | 0.00456581 |
| CYP27A1 | -1.1526164 | 0.03160954 |
| C2 | -1.1527559 | 0.00836781 |
| AGRN | -1.1563289 | 0.02207008 |
| ALDH3B1 | -1.1586766 | 0.02883505 |
| VSTM1 | -1.1602354 | 0.03716233 |
| AC104134.1 | -1.1665129 | 0.04222732 |

| | | |
|---|---|---|
| WDR34 | -1.168414 | 0.00185547 |
| WNT11 | -1.1685384 | 0.04801848 |
| BRCA2 | -1.1703896 | 0.00140632 |
| ITGB8 | -1.1718799 | 0.02467713 |
| MYOF | -1.1726427 | 0.04768915 |
| TMC5 | -1.1736313 | 0.04559516 |
| PTTG1 | -1.1804677 | 0.00284366 |
| LINC00926 | -1.1815218 | 0.0261257 |
| FXYD6 | -1.1830227 | 0.04979261 |
| LRRC25 | -1.1843713 | 0.0224235 |
| ADAMTS14 | -1.1847642 | 0.0271288 |
| PRC1 | -1.185114 | 0.01467928 |
| SMPDL3A | -1.1898012 | 0.0137591 |
| SLCO4A1 | -1.1939711 | 0.01066148 |
| HERC5 | -1.1959207 | 0.00205914 |
| POC1A | -1.1999279 | 0.0052426 |
| CASKIN2 | -1.2000866 | 0.0464278 |
| UBE2T | -1.2020888 | 0.03298769 |
| ARHGAP24 | -1.2024305 | 0.01498328 |
| ORC6 | -1.2028467 | 0.00135042 |
| CD8B2 | -1.2030449 | 0.03092948 |
| SPAG5 | -1.2053226 | 0.0015994 |
| HIST1H2AB | -1.2145609 | 0.00270994 |
| S100A13 | -1.21828 | 0.01706377 |
| PLAUR | -1.2190273 | 0.01658196 |
| CORT | -1.2199966 | 0.02517547 |
| LILRA5 | -1.2222469 | 0.04612604 |
| GPBAR1 | -1.2236619 | 0.01871109 |
| LRRC43 | -1.2260703 | 0.01624546 |
| KIF11 | -1.2266447 | 0.00427217 |
| CATSPER1 | -1.2268093 | 0.04416921 |
| CSF3R | -1.2300455 | 0.04603091 |
| AC007278.2 | -1.2369122 | 0.03561622 |
| RBBP8 | -1.2372224 | 0.00098466 |
| CATSPERD | -1.2408108 | 0.0252231 |
| TYMP | -1.2428584 | 0.01401234 |
| AXL | -1.2437854 | 0.00929944 |
| VDR | -1.2462071 | 0.00989954 |
| FZD2 | -1.2464015 | 0.04896268 |
| C19orf84 | -1.2466293 | 0.01820835 |

| | | |
|---|---|---|
| PLBD1-AS1 | -1.2483937 | 0.02555849 |
| CENPE | -1.2493892 | 0.00134169 |
| CDC42EP4 | -1.2515014 | 0.03304131 |
| CTNNAL1 | -1.2554339 | 0.0069132 |
| THEM5 | -1.257617 | 0.04026518 |
| AC134682.1 | -1.2596776 | 0.03748895 |
| ACPP | -1.261703 | 0.03776994 |
| AC124798.1 | -1.26209 | 0.01936227 |
| SPIB | -1.2625021 | 0.04366908 |
| AP000654.1 | -1.2725542 | 0.01680342 |
| CCNE2 | -1.2742708 | 0.00612309 |
| TMEM255A | -1.27451 | 0.01868925 |
| LIM2 | -1.2767155 | 0.00840464 |
| AC007278.1 | -1.2794149 | 0.03958499 |
| FABP5 | -1.2799274 | 0.00085333 |
| LIF | -1.2806721 | 0.0238548 |
| AL445231.1 | -1.2846013 | 0.03906544 |
| CENPM | -1.2854879 | 0.00063652 |
| AC012507.3 | -1.2876558 | 0.04445663 |
| IGHV5-51 | -1.2908638 | 0.02969223 |
| CENPF | -1.2934649 | 0.00059941 |
| NUF2 | -1.2965158 | 0.00144761 |
| EPOP | -1.2984848 | 0.03115526 |
| NIBAN3 | -1.3006997 | 0.0100732 |
| LMO2 | -1.3029596 | 0.02347776 |
| PILRA | -1.3034391 | 0.01009081 |
| TSHR | -1.3039877 | 0.01287807 |
| PDE2A | -1.3063956 | 0.02078649 |
| AC009570.2 | -1.3064854 | 0.01975228 |
| ADRA2A | -1.3069555 | 0.02449824 |
| OXTR | -1.3092292 | 0.01596912 |
| DGKK | -1.3095848 | 0.03109225 |
| CENPN | -1.3121127 | 0.00431543 |
| IFITM10 | -1.3145571 | 0.03092542 |
| KIF24 | -1.318387 | 0.00849495 |
| TXNDC5 | -1.3186398 | 0.02050268 |
| KLRC1 | -1.31957 | 0.03062784 |
| TMTC1 | -1.3199543 | 0.03675782 |
| FAM20A | -1.3214375 | 0.03460981 |
| RHD | -1.3216942 | 0.04083341 |

| | | |
|---|---|---|
| TNS3 | -1.3240284 | 0.02843135 |
| ASTL | -1.3250979 | 0.04672753 |
| TSPAN13 | -1.3253025 | 0.01820582 |
| ZNF385A | -1.3262224 | 0.02796374 |
| AL359851.1 | -1.3266407 | 0.02483023 |
| HIST1H2AH | -1.3276364 | 0.00104433 |
| CCNB1 | -1.3297106 | 0.00598739 |
| C5AR1 | -1.3316521 | 0.00634868 |
| AC068279.2 | -1.3341313 | 0.00763839 |
| CD86 | -1.3364077 | 0.00506035 |
| CEBPD | -1.3373656 | 0.00265001 |
| FAM30A | -1.3386215 | 0.00649305 |
| MCM2 | -1.3399649 | 0.00206994 |
| AC011472.2 | -1.3418894 | 0.03674246 |
| CD22 | -1.3482428 | 0.03165778 |
| P2RY13 | -1.349327 | 0.04043005 |
| CLEC17A | -1.3499291 | 0.01424607 |
| MSR1 | -1.3560334 | 0.03961022 |
| AURKA | -1.3574679 | 0.02135817 |
| ATP9A | -1.3664522 | 0.02297687 |
| CHEK1 | -1.3691437 | 0.00144449 |
| HIST1H1B | -1.3697948 | 0.00127607 |
| ZWINT | -1.3703687 | 0.00393086 |
| ZNF695 | -1.3722119 | 0.02266155 |
| HOXB5 | -1.3729964 | 0.04359213 |
| PRR29-AS1 | -1.3741416 | 0.03394445 |
| WDR62 | -1.3756219 | 0.00189015 |
| BCAN | -1.3777862 | 0.03671618 |
| AC096667.1 | -1.3793593 | 0.04732903 |
| SLC6A4 | -1.3822837 | 0.02825084 |
| STEAP3 | -1.383108 | 0.04178961 |
| SAPCD2 | -1.383791 | 0.00508869 |
| HMOX1 | -1.3842716 | 0.00197656 |
| CIT | -1.3860725 | 0.00226025 |
| TNFRSF21 | -1.3882186 | 0.01335562 |
| GMNN | -1.3884288 | 0.00171752 |
| CD19 | -1.3899162 | 0.01844868 |
| C2orf66 | -1.3944238 | 0.02406663 |
| TMEM176B | -1.3972037 | 0.04998131 |
| ZBTB32 | -1.4011442 | 0.00408299 |

| | | |
|---|---|---|
| CLEC7A | -1.4014452 | 0.02585995 |
| OPHN1 | -1.4025169 | 0.03339953 |
| AP001610.2 | -1.4031804 | 0.02599282 |
| PDGFC | -1.4036168 | 0.04733483 |
| GNB4 | -1.4070946 | 0.0027117 |
| CD68 | -1.4176204 | 0.01316153 |
| AL353147.1 | -1.4206737 | 0.031753 |
| CD24 | -1.4221902 | 0.01404601 |
| HOXB-AS3 | -1.4245174 | 0.0194924 |
| HIST1H2BM | -1.4267808 | 0.00275935 |
| PLAAT2 | -1.4279502 | 0.02887917 |
| FCGR2A | -1.4284943 | 0.02960941 |
| CPNE5 | -1.4294857 | 0.00639526 |
| IGHV5-10-1 | -1.4298791 | 0.02437539 |
| GPR162 | -1.4318705 | 0.04016573 |
| HAVCR2 | -1.4319834 | 0.0006834 |
| HIST1H2AL | -1.432669 | 0.00095083 |
| KIF7 | -1.4336615 | 0.03229051 |
| TNFAIP2 | -1.4341261 | 0.02941596 |
| DEPDC1 | -1.4344297 | 0.00427227 |
| E2F1 | -1.4348354 | 0.00073541 |
| SYCE2 | -1.4352751 | 0.00848855 |
| HIST1H4I | -1.4381627 | 0.00137404 |
| MEF2C | -1.4385179 | 0.0331385 |
| LINC02785 | -1.4447909 | 0.03188743 |
| CD38 | -1.4459451 | 0.00997303 |
| EME1 | -1.4462958 | 0.03657569 |
| MYO1E | -1.4465321 | 0.00551337 |
| IGKV1-12 | -1.4465743 | 0.0060049 |
| CYP1B1 | -1.4483076 | 0.01925448 |
| SLC27A2 | -1.4505784 | 0.00333337 |
| AC007250.1 | -1.452734 | 0.03195809 |
| POLE2 | -1.4564501 | 0.00289409 |
| JUP | -1.4567775 | 0.00209772 |
| IL13RA1 | -1.4596944 | 0.04103128 |
| DMC1 | -1.4603727 | 0.00317472 |
| LINC02156 | -1.4664235 | 0.02896207 |
| HNMT | -1.4733018 | 0.03689779 |
| MATN4 | -1.4737261 | 0.03415837 |
| LAG3 | -1.4759243 | 0.00391327 |

| | | |
|---|---|---|
| PLA1A | -1.4796603 | 0.04624503 |
| PAX5 | -1.4808133 | 0.01182587 |
| PTX3 | -1.4819218 | 0.04947554 |
| SYCE3 | -1.4826345 | 0.01142077 |
| DSCC1 | -1.4834695 | 0.00271259 |
| U62317.4 | -1.4851495 | 0.00683726 |
| AC004130.2 | -1.4852285 | 0.00638861 |
| SERPINA1 | -1.4856051 | 0.00887579 |
| EXO1 | -1.4869054 | 0.00352465 |
| IL10 | -1.4929052 | 0.03562055 |
| RAB39A | -1.4938631 | 0.01693164 |
| TOP2A | -1.4984022 | 0.00025064 |
| CDCA3 | -1.499831 | 0.00740019 |
| SGO1 | -1.5007278 | 0.00139764 |
| EBI3 | -1.5051795 | 0.01711776 |
| P2RY6 | -1.5087107 | 0.03318778 |
| CKS2 | -1.5095208 | 0.00041374 |
| IFI30 | -1.5100951 | 0.01090918 |
| ISG15 | -1.5102703 | 0.02801064 |
| EBF1 | -1.514538 | 0.01956311 |
| ATP11A-AS1 | -1.5180852 | 0.03884812 |
| HIST1H2BO | -1.5218405 | 0.00255101 |
| MARCKS | -1.5235817 | 0.03538543 |
| OXCT2 | -1.5245842 | 0.04409454 |
| HIST1H2BH | -1.5248719 | 0.00034653 |
| FCRL2 | -1.5251828 | 0.04959895 |
| CABLES1 | -1.529338 | 0.02299555 |
| MNS1 | -1.5383192 | 0.01126172 |
| PXDC1 | -1.5425241 | 0.04136963 |
| MTFR2 | -1.5453571 | 0.00073682 |
| AC026202.2 | -1.5464698 | 0.02300843 |
| TMEM176A | -1.5467752 | 0.0336926 |
| SCN1B | -1.550833 | 0.04786645 |
| CDCA8 | -1.5552594 | 0.00042023 |
| SPI1 | -1.5573512 | 0.02800967 |
| CDKN3 | -1.5593374 | 0.00050869 |
| ZNF804A | -1.5616123 | 0.00465556 |
| AL445985.2 | -1.5640306 | 0.01891987 |
| PLSCR4 | -1.5646875 | 0.01574132 |
| CENPW | -1.5653678 | 0.00026848 |

| | | |
|---|---|---|
| NEK2 | -1.5663946 | 0.00321139 |
| KCNN3 | -1.5705478 | 0.01980536 |
| STIL | -1.5710091 | 0.00076249 |
| SEZ6L | -1.5734946 | 0.03678716 |
| GINS2 | -1.5780604 | 0.00059941 |
| MACC1 | -1.5795442 | 0.00375023 |
| HOXB6 | -1.5825437 | 0.01926954 |
| BTK | -1.5846444 | 0.01724687 |
| CEP55 | -1.5847902 | 0.00095154 |
| PIR | -1.5879156 | 0.00810873 |
| NRP1 | -1.5889583 | 0.0194455 |
| FOSL1 | -1.589551 | 0.04126107 |
| CCNJL | -1.5897758 | 0.01899536 |
| ECM2 | -1.5981689 | 0.01622032 |
| TLR8 | -1.5998523 | 0.02308326 |
| DDO | -1.606907 | 0.04865302 |
| PIMREG | -1.6101645 | 0.00479549 |
| C4B | -1.6139681 | 0.0224162 |
| UBE2R2-AS1 | -1.6143129 | 0.01300616 |
| DEPDC1B | -1.616292 | 0.00071583 |
| SH2B2 | -1.6222234 | 0.0015344 |
| HESX1 | -1.6227228 | 0.03958335 |
| AP001020.2 | -1.6390542 | 0.03582013 |
| PLS3 | -1.6393228 | 0.00839604 |
| BUB1 | -1.6453431 | 0.00159281 |
| CPED1 | -1.6515484 | 0.00467097 |
| IFI44 | -1.6517061 | 0.00403402 |
| PLVAP | -1.6549092 | 0.03489409 |
| TICRR | -1.6564609 | 0.00657918 |
| C17orf53 | -1.6573595 | 0.01714784 |
| ROR2 | -1.6590247 | 0.01953218 |
| NID1 | -1.6605894 | 0.01231802 |
| CENPU | -1.665918 | 0.00191087 |
| SLC22A4 | -1.670305 | 0.01877226 |
| LAMP3 | -1.6713536 | 0.00946584 |
| CLEC4C | -1.6722256 | 0.04872844 |
| IFI6 | -1.6743184 | 0.00790392 |
| ZNF503 | -1.6833664 | 0.03714522 |
| TMEM51 | -1.6838466 | 0.03761586 |
| SYCP2L | -1.6929046 | 0.02761995 |

| | | |
|---|---|---|
| ZNF215 | -1.6946799 | 0.0400733 |
| KHDRBS2 | -1.6981938 | 0.00961698 |
| AL133467.1 | -1.7030969 | 0.00399567 |
| AC017002.3 | -1.7084927 | 0.00585161 |
| IL1B | -1.7098112 | 0.00557733 |
| FAM72B | -1.7194099 | 0.00167179 |
| RDM1 | -1.7242768 | 0.00166879 |
| TYMSOS | -1.7244319 | 0.002263 |
| SPC24 | -1.7254108 | 0.00046098 |
| IL6 | -1.7307843 | 0.02673716 |
| CCDC150 | -1.7320816 | 0.00181079 |
| CLEC10A | -1.7339409 | 0.03058816 |
| S1PR3 | -1.7362071 | 0.0390423 |
| ODF3B | -1.7397902 | 0.00017577 |
| ASMT | -1.7414976 | 0.00153683 |
| HIST1H2AJ | -1.741777 | 0.00066753 |
| SLC6A20 | -1.7419834 | 0.00743792 |
| AL162411.1 | -1.7428887 | 0.04330198 |
| KIF14 | -1.7434309 | 0.00123678 |
| FCN1 | -1.7437727 | 0.04724008 |
| CDKN1A | -1.7519197 | 0.01508961 |
| CXCL2 | -1.7560294 | 0.04577536 |
| TRIP13 | -1.7560793 | 0.00045009 |
| KNL1 | -1.7590961 | 0.00278762 |
| TRIB1 | -1.7662672 | 0.0053502 |
| FCAR | -1.7706224 | 0.03157711 |
| TEDC2 | -1.7713395 | 0.00120715 |
| BUB1B | -1.772303 | 0.00124752 |
| RAD51AP1 | -1.7738719 | 0.00098677 |
| TPX2 | -1.7761791 | 0.00089267 |
| IGF2BP3 | -1.7835214 | 0.02149757 |
| MX1 | -1.7859772 | 0.02493838 |
| HMGB3 | -1.7895692 | 0.00542272 |
| PID1 | -1.7940951 | 0.03251405 |
| POLQ | -1.7960423 | 0.00074384 |
| JDP2 | -1.7983833 | 0.02674349 |
| CMPK2 | -1.8000139 | 0.01521834 |
| ORC1 | -1.8050958 | 0.00072322 |
| IL21 | -1.8081006 | 0.00351308 |
| ASF1B | -1.8099291 | 0.00039718 |

| | | |
|---|---|---|
| LOXHD1 | -1.8109801 | 0.02223946 |
| CLIC4 | -1.8143692 | 0.00963274 |
| TTK | -1.8173884 | 0.00113735 |
| NCAPH | -1.8179241 | 0.00114268 |
| HASPIN | -1.8182064 | 0.00093039 |
| PDK4 | -1.8202462 | 0.00637669 |
| KIF18B | -1.8237588 | 0.00177067 |
| SERPINE1 | -1.8244343 | 0.00581795 |
| TGFB2 | -1.8268839 | 0.00601318 |
| TRPM4 | -1.8286621 | 0.00322608 |
| HTR4 | -1.8345816 | 0.04571007 |
| KIF17 | -1.8442617 | 0.03645968 |
| CNTNAP2 | -1.8445734 | 0.01360262 |
| RAD51 | -1.8450304 | 0.00062109 |
| OASL | -1.8628999 | 0.00128817 |
| CDC25C | -1.8631766 | 0.0032812 |
| ESCO2 | -1.8652188 | 0.00061239 |
| PARM1 | -1.8692652 | 0.00491005 |
| NUSAP1 | -1.8741654 | 0.00186184 |
| PRRG4 | -1.8745016 | 0.00123242 |
| CD180 | -1.8752318 | 0.00291637 |
| FAM72D | -1.8802334 | 0.01512128 |
| RAD54L | -1.8856507 | 0.00085195 |
| ESPL1 | -1.8894236 | 0.00154239 |
| KIFC1 | -1.9088267 | 0.00017205 |
| FBXO43 | -1.9099757 | 0.01275993 |
| CLSPN | -1.9113635 | 0.00010056 |
| KRT81 | -1.9127595 | 0.02512414 |
| HOXB9 | -1.9136928 | 0.0147023 |
| AC009961.4 | -1.9196002 | 0.03208066 |
| AC099524.1 | -1.921942 | 0.00458887 |
| FGF2 | -1.9243498 | 0.00691436 |
| NECTIN2 | -1.9275033 | 0.00524443 |
| MIXL1 | -1.9364134 | 0.00865325 |
| KIF15 | -1.936731 | 0.00076598 |
| AL133215.2 | -1.9385862 | 0.00178178 |
| LINC01151 | -1.9465146 | 0.02113627 |
| ADM | -1.9480038 | 0.00633284 |
| ASPM | -1.9481309 | 0.00296086 |
| CDC25A | -1.9536461 | 0.00071713 |

| | | |
|---|---|---|
| DENND5B | -1.9536911 | 0.00394901 |
| CDT1 | -1.9554085 | 0.00017654 |
| HIST1H3F | -1.9579008 | 0.00022093 |
| GMPR | -1.9583738 | 0.00457674 |
| KIF20A | -1.959653 | 0.00085822 |
| ERICH3 | -1.9611617 | 0.00130192 |
| NCAPG | -1.964543 | 0.00024067 |
| CXCR6 | -1.9657555 | 0.00028493 |
| DGKG | -1.9780964 | 0.03674618 |
| UHRF1 | -1.9800998 | 0.00017131 |
| ERCC6L | -1.983733 | 0.00068237 |
| ADRA2B | -1.9859322 | 0.04946976 |
| AC068989.1 | -1.9891349 | 0.00600742 |
| PLD4 | -1.9905835 | 0.04167566 |
| FBXO39 | -1.9930825 | 0.01034522 |
| RAB38 | -1.9944674 | 0.00108034 |
| TROAP | -1.999986 | 0.00162124 |
| DLGAP5 | -2.0032042 | 0.00059056 |
| MELK | -2.0160559 | 0.00048924 |
| PCYT1B | -2.020095 | 0.0155804 |
| AL590666.2 | -2.0226904 | 0.00604535 |
| CDC20 | -2.0260237 | 0.00016143 |
| RHOBTB1 | -2.0343378 | 0.02148027 |
| HIST1H3C | -2.0348774 | 0.00013476 |
| TUBA8 | -2.0358735 | 0.04522916 |
| LAMP5 | -2.0385803 | 0.02038508 |
| AL513493.1 | -2.0416975 | 0.04482131 |
| CDC6 | -2.0489595 | 0.00017323 |
| SLC6A12 | -2.0531302 | 0.0196124 |
| PRICKLE1 | -2.0537531 | 0.00637756 |
| PANX2 | -2.0571616 | 0.02796221 |
| FOXM1 | -2.0576547 | 0.00092186 |
| AC007381.1 | -2.0604858 | 0.04971735 |
| UCHL1 | -2.0628872 | 0.00144537 |
| FFAR2 | -2.0645523 | 0.00596448 |
| HRK | -2.0650121 | 0.01353864 |
| AL353152.2 | -2.0677538 | 0.013603 |
| PLK1 | -2.0692927 | 0.00074808 |
| TNS1 | -2.0753259 | 0.02565429 |
| CDC45 | -2.0762386 | 0.0001307 |

| | | |
|---|---|---|
| AUNIP | -2.0816547 | 0.00111074 |
| IL1RN | -2.0862431 | 0.04374952 |
| HMMR | -2.0880784 | 0.0004888 |
| IQGAP3 | -2.0884383 | 0.00767786 |
| AC138207.9 | -2.1091798 | 0.04402662 |
| CXCL11 | -2.1101291 | 0.04498424 |
| SLC8A3 | -2.1152633 | 0.03734835 |
| SKA1 | -2.1166763 | 0.00141675 |
| TMEM171 | -2.1177005 | 0.01209739 |
| CXCL8 | -2.1198668 | 0.02632346 |
| DTL | -2.1261502 | 0.00023946 |
| LINC02397 | -2.1282083 | 0.00763401 |
| KIF4A | -2.1327545 | 0.0012331 |
| AC243964.3 | -2.1332022 | 0.03405311 |
| IL27 | -2.1363915 | 0.04297532 |
| IGKV1D-33 | -2.1368323 | 0.0252916 |
| BLNK | -2.1371376 | 0.00505111 |
| DIAPH3 | -2.1403787 | 0.00348605 |
| CKAP2L | -2.1409789 | 0.00764577 |
| KIF2C | -2.142267 | 0.00024682 |
| CDCA2 | -2.1519106 | 0.00034155 |
| AC104699.1 | -2.1570637 | 0.00372045 |
| ILDR1 | -2.1589856 | 0.00731505 |
| MEIS1-AS2 | -2.1634447 | 0.03988706 |
| AL591115.1 | -2.163982 | 0.02698665 |
| GHSR | -2.1652817 | 0.0191786 |
| XK | -2.1657353 | 0.02737262 |
| FCGR1B | -2.1693158 | 0.03522338 |
| E2F7 | -2.1731558 | 0.00542007 |
| AC093159.1 | -2.1768932 | 0.02004644 |
| SHCBP1 | -2.1795317 | 0.00066262 |
| MAFB | -2.1899888 | 0.00580841 |
| CDK1 | -2.193362 | 0.00010571 |
| MND1 | -2.1975137 | 0.00042548 |
| FBXO16 | -2.199541 | 0.01550138 |
| FAM111B | -2.2115826 | 0.00048894 |
| NEXN-AS1 | -2.211711 | 0.03455941 |
| HCAR2 | -2.2133011 | 0.01353895 |
| MYEOV | -2.2134021 | 0.01655351 |
| APOBEC3B | -2.2262453 | 0.00037008 |

| | | |
|---|---|---|
| DERL3 | -2.2294126 | 0.00069422 |
| THBS1 | -2.2325459 | 0.03186494 |
| SKA3 | -2.2369148 | 0.00128728 |
| GLDC | -2.2376558 | 0.00088789 |
| NRIR | -2.2384693 | 0.01660214 |
| TK1 | -2.2426085 | 0.00012275 |
| CAV1 | -2.2449666 | 0.04009824 |
| PNOC | -2.24699 | 0.00220361 |
| CENPA | -2.2485743 | 0.001531 |
| CRYBB1 | -2.2554442 | 0.03320523 |
| COBLL1 | -2.2572936 | 0.00140649 |
| MKI67 | -2.2621094 | 0.00323593 |
| AL031005.1 | -2.2624811 | 0.0477074 |
| FPR3 | -2.2650051 | 0.00250743 |
| IGF1 | -2.2657506 | 0.00296577 |
| SUCNR1 | -2.2748331 | 0.04659084 |
| LILRB4 | -2.2771683 | 0.01281613 |
| AL138720.1 | -2.2819315 | 0.00850133 |
| HIST1H3B | -2.2845591 | 0.00253155 |
| IGKC | -2.2953849 | 0.00455859 |
| AURKB | -2.2995639 | 0.00025235 |
| IGKV3D-15 | -2.3019443 | 0.00139182 |
| GNG4 | -2.3113965 | 0.00308879 |
| MT1G | -2.3166729 | 0.00819236 |
| F2RL3 | -2.3187328 | 0.0297549 |
| SPC25 | -2.3196806 | 0.00104926 |
| SLC2A5 | -2.3282721 | 0.01147885 |
| PTCRA | -2.3284778 | 0.04171725 |
| SDS | -2.331196 | 0.00989964 |
| PACSIN1 | -2.3411393 | 0.00059723 |
| AL158168.1 | -2.3565617 | 0.00670494 |
| POU2AF1 | -2.3574163 | 0.00017463 |
| RNF152 | -2.3643148 | 0.02807399 |
| TRAPPC3L | -2.37455 | 0.04731007 |
| FCRL5 | -2.3840068 | 0.00304355 |
| PBK | -2.384612 | 0.00097549 |
| CCNB2 | -2.387475 | 0.00156685 |
| PTCHD1 | -2.3891936 | 0.01617829 |
| PKMYT1 | -2.3899501 | 0.00178988 |
| MIR503HG | -2.3951371 | 0.01091468 |

| | | |
|---|---|---|
| MCM10 | -2.3988403 | 0.00049899 |
| ACTG2 | -2.3989747 | 0.00713394 |
| CCNA2 | -2.4005834 | 0.00024508 |
| TMEM158 | -2.4102129 | 0.04691486 |
| IGHV3-49 | -2.4139426 | 0.02395127 |
| GAB1 | -2.4180208 | 0.01834194 |
| TNFRSF13B | -2.4195293 | 0.00085458 |
| ANLN | -2.420077 | 0.00644085 |
| XIRP2 | -2.4211147 | 0.0176649 |
| RAPGEF5 | -2.4237995 | 0.00851624 |
| GPRC5D | -2.4359723 | 0.0027894 |
| SMOX | -2.4433469 | 0.02112748 |
| IGKV2-28 | -2.4442039 | 0.01782883 |
| RRM2 | -2.4595308 | 0.00035681 |
| PCLAF | -2.460893 | 0.00049896 |
| KIF23 | -2.4671796 | 0.00939429 |
| LINC00853 | -2.470423 | 0.02719672 |
| SPACA3 | -2.4738287 | 0.03893696 |
| SLFN14 | -2.4762607 | 0.02295015 |
| E2F8 | -2.4831464 | 0.0031207 |
| TYMS | -2.4857453 | 0.00092201 |
| TIMD4 | -2.4901904 | 0.00016505 |
| IGHV1-17 | -2.4911625 | 0.006356 |
| IGKV3OR2-268 | -2.4933082 | 0.00063373 |
| MOXD1 | -2.5117543 | 0.00400673 |
| CDCA5 | -2.5149358 | 0.00122439 |
| GALNT14 | -2.5183416 | 0.0016455 |
| GTSE1 | -2.5212093 | 0.00248283 |
| AC105415.1 | -2.5409815 | 0.04076577 |
| FAM81B | -2.5418795 | 0.0327053 |
| BIRC5 | -2.5514984 | 0.00639374 |
| PKHD1L1 | -2.557163 | 0.00643556 |
| LINC00189 | -2.5595274 | 0.01206085 |
| VEPH1 | -2.5708618 | 0.02153526 |
| SPAG6 | -2.5890745 | 0.0283477 |
| IFIT1 | -2.5923194 | 0.00612432 |
| CRYM | -2.6117452 | 0.00674163 |
| EPHB2 | -2.6200871 | 0.01249286 |
| HIST1H3G | -2.6256832 | 0.00588472 |
| AC104024.1 | -2.6431261 | 0.00888529 |

| | | |
|---|---|---|
| SERPING1 | -2.6741946 | 0.00025455 |
| IGKV3D-20 | -2.6812828 | 0.00115624 |
| INKA2-AS1 | -2.6888098 | 0.04654533 |
| BHLHA15 | -2.6979163 | 0.00744864 |
| PSD2 | -2.7092796 | 0.00382745 |
| SPDYC | -2.7199365 | 0.02130422 |
| IGKV1D-8 | -2.7302381 | 0.01740539 |
| OTOF | -2.7354567 | 0.02013755 |
| HJURP | -2.7381962 | 0.00620894 |
| UBE2C | -2.7384956 | 0.0109262 |
| AC116903.2 | -2.7417974 | 0.01170547 |
| MYBL2 | -2.7524473 | 0.00154938 |
| CAV2 | -2.7650414 | 0.01000399 |
| USP41 | -2.7747105 | 0.03220939 |
| AC037198.1 | -2.7826023 | 0.00202737 |
| EPHA2 | -2.7937765 | 0.03769463 |
| AC008870.3 | -2.8114249 | 0.04137845 |
| ANKRD45 | -2.8143992 | 0.00112823 |
| ARMC3 | -2.8251788 | 0.02169741 |
| BHLHE41 | -2.8427727 | 0.00035747 |
| CXCL13 | -2.8606586 | 0.01002522 |
| LINC02576 | -2.8780646 | 0.00336896 |
| IFIT3 | -2.8906396 | 0.00017488 |
| AP001189.3 | -2.890818 | 0.02003316 |
| LINC01501 | -2.9105185 | 0.01032749 |
| BMP6 | -2.92566 | 0.00084384 |
| LAMC1 | -2.926655 | 0.00298225 |
| LINC00487 | -2.9303088 | 0.00202654 |
| LIPH | -2.9373194 | 0.01555043 |
| EGF | -2.9625174 | 0.03414075 |
| CHAD | -2.9776055 | 0.00401662 |
| PDE3A | -2.9867138 | 0.01484259 |
| CCL2 | -2.988947 | 0.00784592 |
| AC012236.1 | -3.0044287 | 0.00030641 |
| P2RY12 | -3.0166556 | 0.04049677 |
| AC105094.2 | -3.0214601 | 0.02488726 |
| IGHG2 | -3.0416251 | 0.01134504 |
| AL133405.2 | -3.0421885 | 0.00306369 |
| AC025259.3 | -3.0570228 | 0.02966178 |
| AC017002.6 | -3.0792192 | 0.01282861 |

| | | |
|---|---|---|
| C1QA | -3.1005426 | 0.00046878 |
| AL954642.1 | -3.1305038 | 0.02979505 |
| ABLIM3 | -3.1706775 | 0.00627254 |
| LINC02757 | -3.174614 | 0.01365465 |
| TNFRSF17 | -3.1807301 | 0.00044811 |
| IFI44L | -3.184837 | 0.0017076 |
| IGHJ4 | -3.2129491 | 0.01819524 |
| IGHV4-59 | -3.2250219 | 0.03446061 |
| CCR1 | -3.2317342 | 0.00133166 |
| MZB1 | -3.2379506 | 0.00016656 |
| AVPR1A | -3.2764708 | 0.01582276 |
| AL445426.1 | -3.2815169 | 0.01775245 |
| IGLV2-11 | -3.301069 | 0.00938784 |
| IGKV2D-28 | -3.3057307 | 0.04199546 |
| LCN2 | -3.3477607 | 0.02033291 |
| AF127936.1 | -3.3672669 | 0.00168969 |
| LINC01857 | -3.3789445 | 0.00409119 |
| IGHV2-5 | -3.3818661 | 0.00101296 |
| BVES | -3.39666 | 0.02590275 |
| LURAP1L | -3.4264296 | 0.00867407 |
| IGLV3-10 | -3.4288382 | 0.04333219 |
| AC020914.1 | -3.432926 | 0.01471729 |
| IGLJ1 | -3.4533729 | 0.00180066 |
| AL451164.1 | -3.4674908 | 0.01715131 |
| AP002478.1 | -3.4678743 | 0.00561216 |
| PDGFRA | -3.4735014 | 0.01051219 |
| IGLV6-57 | -3.4898698 | 0.00013791 |
| IGHV1-46 | -3.599833 | 0.00015055 |
| SH3TC2 | -3.6162758 | 0.00521478 |
| IGHD | -3.6248193 | 0.03269471 |
| IGHV1-2 | -3.6275588 | 0.00013761 |
| IGHGP | -3.6380717 | 0.00020241 |
| IGLVI-70 | -3.7073696 | 0.00113786 |
| IGLV1-44 | -3.7126303 | 2.00E-05 |
| IGLV1-41 | -3.7132646 | 0.00114351 |
| ITGA8 | -3.7156375 | 0.02187117 |
| CHST8 | -3.7599861 | 0.02260129 |
| IGHV3-30 | -3.7707072 | 5.68E-05 |
| IGKV2D-30 | -3.798096 | 0.00547455 |
| IGLC2 | -3.806462 | 0.00918297 |

| | | |
|---|---|---|
| JCHAIN | -3.8145874 | 0.00272246 |
| IGKV1D-39 | -3.8364515 | 0.01039396 |
| C1QB | -3.8414748 | 0.00189517 |
| IGHV1-3 | -3.8739375 | 0.00988856 |
| IGLV2-14 | -3.9445043 | 0.04048158 |
| APOBEC3A | -3.9568453 | 0.00924825 |
| SDC1 | -3.9948913 | 0.00502536 |
| IGKV2-30 | -4.2381799 | 0.00030436 |
| AC114752.2 | -4.2648517 | 0.00489236 |
| CALHM5 | -4.2969957 | 0.01659587 |
| IGKV1-39 | -4.333104 | 0.0006654 |
| IGHM | -4.3401276 | 0.00569589 |
| KITLG | -4.3989066 | 0.01337823 |
| IGHV4-39 | -4.409078 | 0.02319665 |
| LINC01484 | -4.4187443 | 0.00886639 |
| IGLV7-46 | -4.4345687 | 2.56E-05 |
| SERPINB2 | -4.4936879 | 0.0040804 |
| IGLV3-19 | -4.5102906 | 0.00028351 |
| IGLV4-60 | -4.5511129 | 0.00044577 |
| IGLC7 | -4.5794744 | 0.00842697 |
| KCTD14 | -4.6972673 | 0.00512814 |
| IGKV1-27 | -4.702591 | 0.00160261 |
| IGLC1 | -4.7040112 | 0.01112846 |
| IGHV2-26 | -4.723562 | 0.00788573 |
| IGHG1 | -4.7367185 | 0.0012837 |
| C1QC | -4.7982021 | 0.00020847 |
| IGHV3-15 | -4.8097718 | 0.0003704 |
| AC135068.2 | -4.8238198 | 0.03523781 |
| GPR84 | -4.8376706 | 0.00117578 |
| IGHV4-28 | -4.8908302 | 0.00026996 |
| BHLHE23 | -4.8963613 | 0.00805177 |
| IGKV1D-27 | -4.9446217 | 0.03016038 |
| IFI27 | -5.0242725 | 0.01486008 |
| AC025887.2 | -5.0778987 | 0.02500981 |
| SIGLEC1 | -5.1720633 | 0.00012515 |
| IGKV6-21 | -5.1842014 | 0.00187448 |
| IGKV3-20 | -5.2646761 | 2.95E-05 |
| IGKV3D-11 | -5.3391259 | 0.00062843 |
| IGKV3-15 | -5.5543052 | 0.00135117 |
| CXCL10 | -5.600364 | 0.00299073 |

| | | |
|---|---|---|
| IGKV2D-24 | -5.678688 | 0.00698591 |
| KLHL14 | -5.6969898 | 2.72E-05 |
| IGLV3-25 | -5.7221726 | 0.00997444 |
| IGLC3 | -5.744436 | 0.00059204 |
| IGKV1D-43 | -5.7559155 | 0.01531733 |
| IGLV3-6 | -5.8522409 | 0.01545823 |
| IGHV1-24 | -5.888559 | 0.00059445 |
| IGKV1D-37 | -5.9583773 | 0.01432537 |
| IGKV3-11 | -5.9781898 | 0.03078403 |
| IGHV6-1 | -5.978317 | 4.36E-06 |
| IGLV3-16 | -6.0035861 | 0.0005358 |
| IGLV5-37 | -6.0349078 | 0.00016214 |
| IGHV4-4 | -6.0633453 | 0.0004444 |
| IGKV1D-16 | -6.1007878 | 0.0092931 |
| IGKV2D-26 | -6.1115656 | 0.01239061 |
| IGHV3OR16-9 | -6.2028057 | 0.0126324 |
| SCGB1C1 | -6.3127685 | 0.0108568 |
| IGHV3-64 | -6.3506713 | 0.03306215 |
| IGKV7-3 | -6.3880159 | 0.01076587 |
| IGHV4OR15-8 | -6.6172604 | 0.00927616 |
| IGKV5-2 | -6.7472306 | 0.00931995 |
| IGHV3-23 | -6.7884469 | 1.20E-05 |
| IGHV4-55 | -6.8424366 | 7.86E-05 |
| IGLV3-21 | -6.8747268 | 0.02710329 |
| IGHV1-58 | -6.9535285 | 0.00268954 |
| IGLV7-43 | -7.018437 | 2.15E-05 |
| IGLV5-45 | -7.2456816 | 0.0016337 |
| IGHV3-22 | -7.3682283 | 0.00516594 |
| IGHV3-20 | -7.3691496 | 5.78E-05 |
| IGLV1-40 | -7.9915251 | 7.76E-06 |
| IGLV9-49 | -8.2271144 | 9.38E-05 |
| IGKV6D-21 | -8.3086348 | 0.00358628 |
| IGHV3-72 | -8.7357577 | 0.00022792 |
| IGKV1-9 | -8.8011474 | 0.01217045 |
| IGLV4-69 | -9.0500897 | 6.46E-05 |
| IGKV1-17 | -9.0527453 | 0.00116231 |
| IGHV1-69D | -9.1513454 | 0.0005238 |
| IGLV10-54 | -9.2981013 | 0.00159694 |

**Table S6:** Differentially expressed genes in paired samples of CD4+ T cells from during viremia vs. ART suppression (n=3) identified using Wilcoxon Rank-sum test (adjusted p-value < 0.05 and $\log_2$(fold-change) >0.25)

| gene | $Log_2$ fold-change | adjusted p-value |
|---|---|---|
| IFI44L | 0.68711536 | 0 |
| IFI6 | 0.61774893 | 0 |
| LY6E | 0.59441547 | 0 |
| ISG15 | 0.57655445 | 0 |
| IFITM1 | 0.57518807 | 0 |
| GNLY | 0.55908517 | 0 |
| MX1 | 0.52526018 | 0 |
| IGLV2-14 | 0.50209092 | 0 |
| XAF1 | 0.49385787 | 0 |
| H3F3B | 0.47568533 | 0 |
| GZMB | 0.47159627 | 5.47E-254 |
| PRF1 | 0.46692619 | 1.69E-297 |
| KLF6 | 0.40884327 | 0 |
| MT2A | 0.40150307 | 0 |
| H1FX | 0.3983851 | 0 |
| IL7R | 0.35222641 | 2.78E-278 |
| NEAT1 | 0.3458239 | 0 |
| EIF2AK2 | 0.32949404 | 0 |
| JCHAIN | 0.32635971 | 0 |
| SP100 | 0.32583939 | 0 |
| EPSTI1 | 0.3213101 | 0 |
| TYMP | 0.31982859 | 0 |
| PTPRCAP | 0.31893667 | 0 |
| TRIM22 | 0.30912806 | 0 |
| NFKBIA | 0.30806242 | 4.87E-287 |
| SAT1 | 0.3076522 | 0 |
| IFITM2 | 0.3064805 | 0 |
| PIK3R1 | 0.30601626 | 0 |
| FTH1 | 0.30502518 | 0 |
| CXCR4 | 0.30326592 | 0 |
| ISG20 | 0.29822877 | 0 |
| PDE4D | 0.29628865 | 0 |
| RORA | 0.29536728 | 0 |
| ZFP36 | 0.29369697 | 0 |
| RGCC | 0.28968505 | 2.34E-29 |
| JUND | 0.28745438 | 5.43E-218 |
| ARL4C | 0.28007898 | 0 |
| ZFP36L2 | 0.27494805 | 0 |
| IRF7 | 0.27451789 | 0 |

| | | |
|---|---|---|
| UBC | 0.27429197 | 0 |
| EML4 | 0.2727441 | 0 |
| FOS | 0.2688643 | 2.96E-103 |
| EZR | 0.26822532 | 0 |
| RNF213 | 0.26642503 | 0 |
| DUSP2 | 0.26544266 | 4.86E-210 |
| PIM3 | 0.26337664 | 1.68E-169 |
| IFITM3 | 0.26018049 | 2.42E-99 |
| BST2 | 0.25578401 | 0 |
| DDX24 | 0.2553014 | 0 |
| PRRC2C | 0.25243728 | 0 |
| LINC00861 | -0.2536066 | 0 |
| RPS18 | -0.2542308 | 0 |
| RPL26 | -0.2557391 | 0 |
| RPL35A | -0.2567487 | 0 |
| MT-ND4 | -0.2603619 | 0 |
| RPS9 | -0.26176 | 0 |
| UCP2 | -0.2650457 | 0 |
| RPL18 | -0.2650766 | 0 |
| RPS13 | -0.2695542 | 0 |
| RACK1 | -0.2730838 | 0 |
| MT-CO3 | -0.2776546 | 0 |
| RPS15A | -0.2867341 | 0 |
| RPS3 | -0.2883389 | 0 |
| RPS12 | -0.2889755 | 0 |
| RPL13 | -0.2996178 | 0 |
| RPS23 | -0.3025982 | 0 |
| EIF3L | -0.3100246 | 0 |
| RPS4X | -0.3121432 | 0 |
| ACTG1 | -0.3132663 | 0 |
| RPS2 | -0.3143116 | 0 |
| RPL10A | -0.3250765 | 0 |
| EEF2 | -0.3317496 | 0 |
| RPS27A | -0.3358043 | 0 |
| MT-CYB | -0.338582 | 0 |
| RPS14 | -0.3423297 | 0 |
| RPS6 | -0.3476099 | 0 |
| RPL10 | -0.3564941 | 0 |
| MT-CO1 | -0.3601173 | 0 |
| RPL3 | -0.3606861 | 0 |
| TCF7 | -0.3618587 | 0 |
| LEF1 | -0.363168 | 0 |
| TPT1 | -0.3642558 | 0 |
| RPS10 | -0.377689 | 0 |
| AL138963.4 | -0.3851407 | 0 |
| RPL5 | -0.39527 | 0 |
| EEF1A1 | -0.3970105 | 0 |
| RPL21 | -0.3998752 | 0 |

| | | |
|---|---|---|
| EEF1G | -0.4082308 | 0 |
| RPS8 | -0.4142539 | 0 |
| RPL4 | -0.4146685 | 0 |
| RPS3A | -0.4364537 | 0 |
| RPS5 | -0.4401782 | 0 |
| EEF1B2 | -0.4613363 | 0 |

**Table S7:** Differentially expressed genes in paired samples of HIV RNA+ CD4+ T cells from during viremia vs. ART suppression (n=3) determined by genewise quasi F-tests on the samples' pseudo-bulked gene counts (p-value < 0.05 and $log_2$(fold-change) >0.25)

| gene | $Log_2$ fold-change | p-value |
|---|---|---|
| IFI27 | 10.6976471 | 6.68E-05 |
| HIV | 2.24138688 | 0.01020525 |
| HIST1H4C | 2.23140196 | 0.00154144 |
| ISG15 | 1.90572946 | 0.01535394 |
| MX1 | 1.78775966 | 0.0361383 |
| SP100 | 1.6738723 | 0.005092 |
| TXNL4A | 1.51095089 | 0.02428004 |
| LYAR | 1.38429987 | 0.0300768 |
| MGAT4A | 1.30751788 | 0.04605694 |
| DUSP2 | 1.28917119 | 0.02037074 |
| CLEC2B | 1.27620858 | 0.00616252 |
| UBE2V1 | 1.24940412 | 0.0444886 |
| TUBB | 1.22790295 | 0.03406084 |
| EID1 | 1.19325251 | 0.01974149 |
| HMGN1 | 1.00084244 | 0.03906102 |
| SRRM1 | 0.93400711 | 0.03373428 |
| SET | 0.92043964 | 0.02650138 |
| CHCHD2 | 0.7804552 | 0.03237411 |
| H3F3B | 0.72190604 | 0.04865253 |
| ITGB1 | -0.7557273 | 0.02683426 |
| YPEL3 | -0.7769544 | 0.04533424 |
| RBL2 | -0.912846 | 0.04692392 |
| IRF1 | -1.1187375 | 0.01971127 |
| RPS10 | -1.3370657 | 0.02406715 |
| SLC38A2 | -1.6941735 | 0.03822155 |

**SUPPLEMENTARY FIGURES**

Figure S1

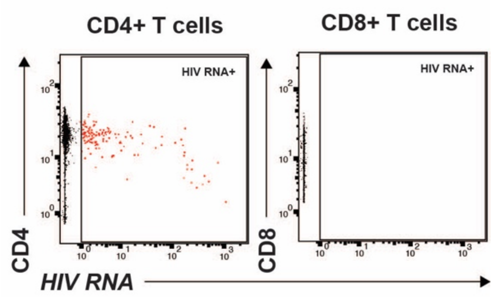

**Figure S2**

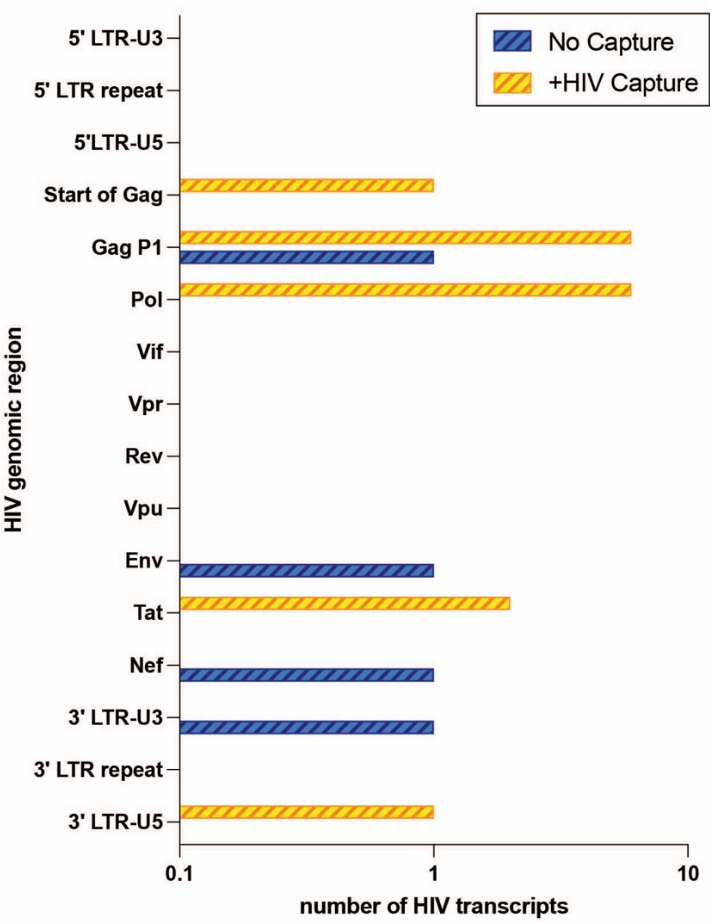

No Capture
+HIV Capture
5' LTR-U3
5' LTR repeat
5'LTR-U5
Start of Gag
Gag P1
Pol
Vif
Vpr
Rev
Vpu
Env
Tat
Nef
3' LTR-U3
3' LTR repeat
3' LTR-U5
HIV genomic region
0.1
1
10
number of HIV transcripts

**Figure S3**

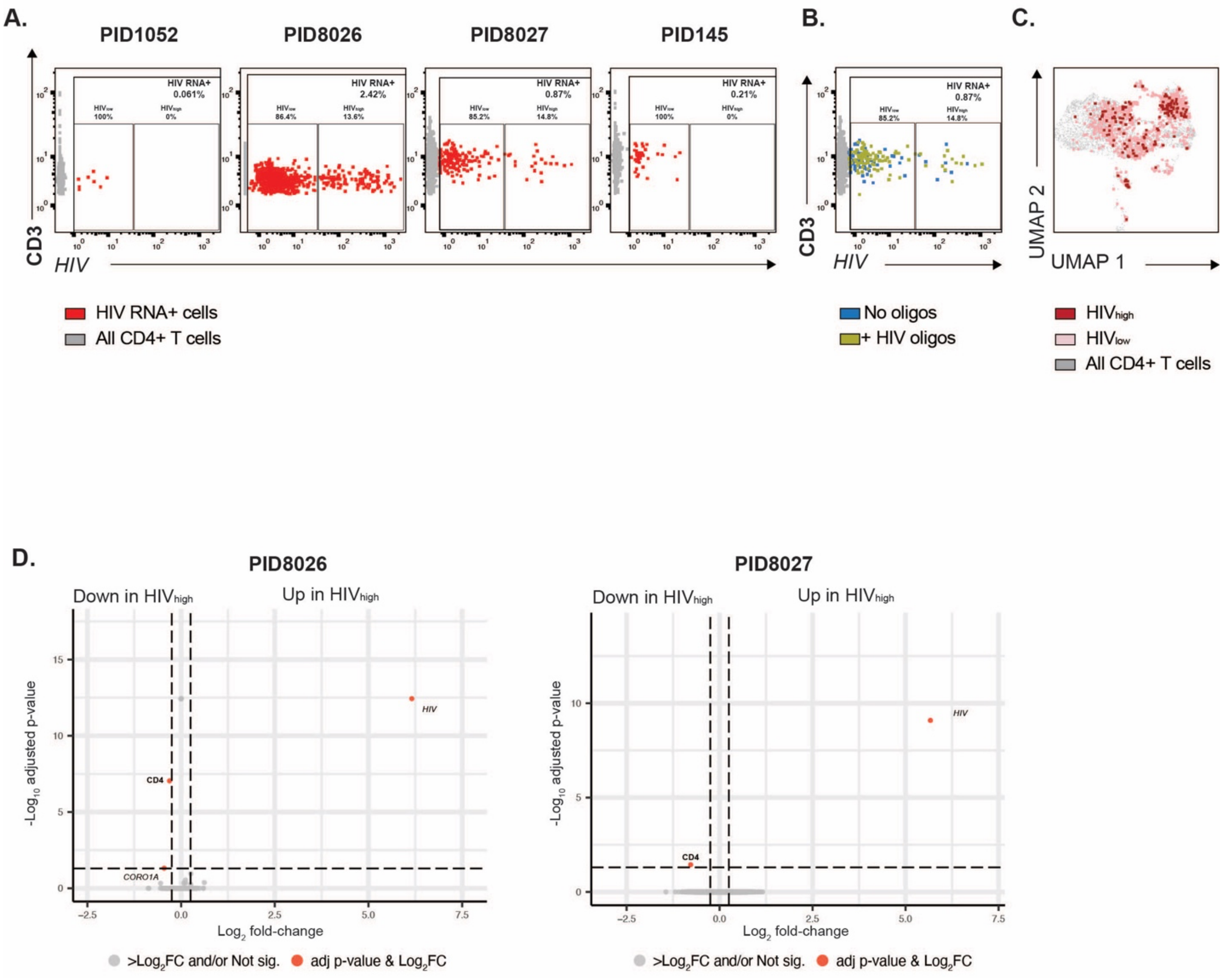
A.
PID1052
PID8026
PID8027
PID145
HIV RNA+ 0.061%
HIV RNA+ 2.42%
HIV RNA+ 0.87%
HIV RNA+ 0.21%
CD3
HIV
HIV RNA+ cells
All CD4+ T cells
B.
HIV RNA+ 0.87%
CD3
HIV
No oligos
+ HIV oligos
C.
UMAP 2
UMAP 1
HIVhigh
HIVlow
All CD4+ T cells
D.
PID8026
PID8027
Down in HIVhigh
Up in HIVhigh
-Log10 adjusted p-value
Log2 fold-change
HIV
CD4
CORO1A
>Log2FC and/or Not sig.
adj p-value & Log2FC

**Figure S4**

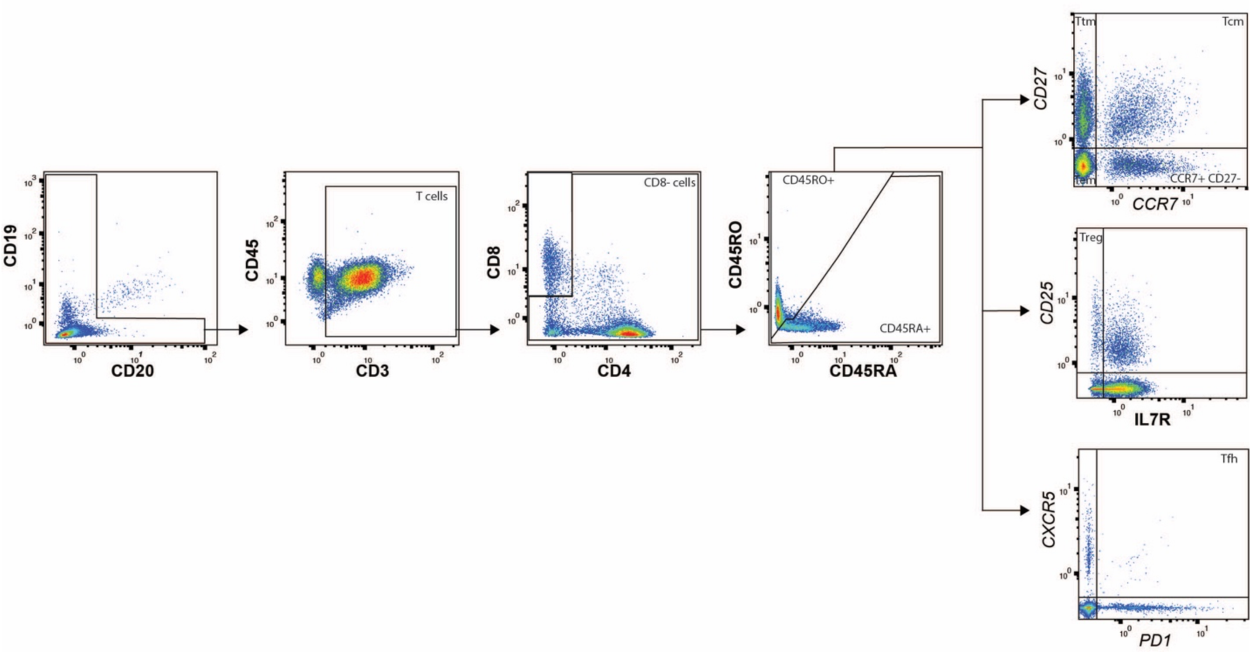
CD19
CD20
CD45
CD3
T cells
CD8
CD4
CD8- cells
CD45RO
CD45RA
CD45RO+
CD45RA+
CD27
CCR7
Ttm
Tcm
Tem
CCR7+ CD27-
CD25
IL7R
Treg
CXCR5
PD1
Tfh

**Figure S5**

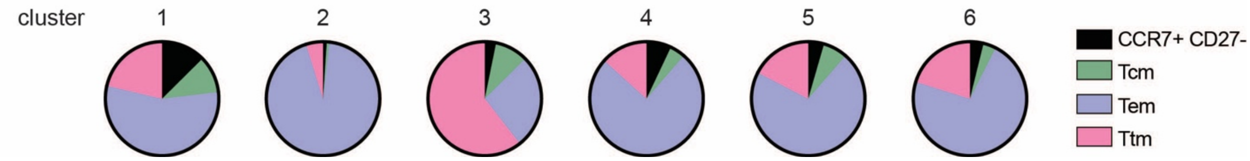

**Figure S6**

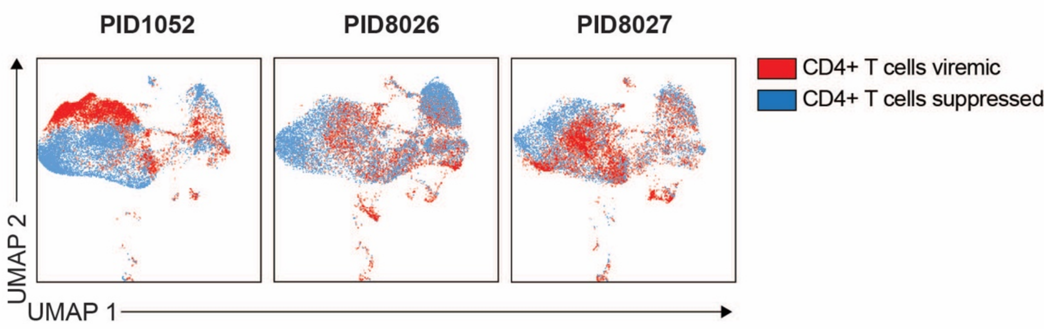
PID1052
PID8026
PID8027
CD4+ T cells viremic
CD4+ T cells suppressed
UMAP 2
UMAP 1

#### Figure S7

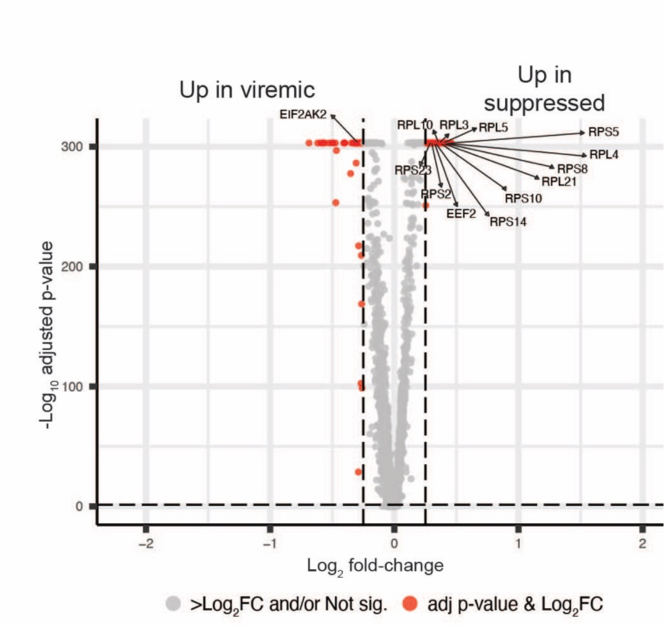

Up in viremic
Up in suppressed
EIF2AK2
RPL10
RPL3
RPL5
RPS5
RPL4
RPS8
RPL21
RPS10
RPS14
EEF2
RPS2
RPS23
300
200
100
0
-Log10 adjusted p-value
−2
−1
0
1
2
Log2 fold-change
>Log2FC and/or Not sig.
adj p-value & Log2FC

#### SUPPLEMENTARY FIGURES LEGENDS

**Figure S1. HIV RNA+ cells are identified among CD4+ but not CD8+ T cells from viremic PWH analyzed by HIV-seq.** CD45+CD3+ T cells analyzed by HIV-seq were gated for CD8- cells (left, to account for potential CD4+ T cells that may have downregulated cell-surface CD4) and CD8+ cells (right), and visualized for cell-surface CD4 and CD8 expression as indicated along with levels of HIV RNA transcripts. Shown are representative results from one (PID8027) of four viremic donors.

**Figure S2. HIV-seq on cells from ART-suppressed PWH increases detection of transcripts mapping to the Gag and Pol regions of HIV-1.** HIV reads from PID1052 and PID8027 during ART suppression were aligned to the HIV-1 subtype B consensus reference genome. The y-axis depicts individual HIV genes, and the x-axis shows the number of detected HIV transcripts, in the absence (blue hatched) vs. presence (yellow hatched) of the HIV capture sequences.

**Figure S3. HIV RNA+ cells expressing low vs. high numbers of HIV transcripts exhibit similar global gene expression profiles but differ in cell-surface CD4 expression.** **A.** HIV RNA+ CD4+ T cells from viremic PWH can be separated into two groups: those harboring 0 to 50 HIV transcripts ($HIV_{low}$), and those with more than 50 HIV transcripts ($HIV_{high}$). HIV RNA+ cells are depicted in red and HIV RNA- cells are depicted in gray. Percentages of total HIV RNA+ cells are indicated in the upper right of each plot, and percentages of $HIV_{low}$ and $HIV_{high}$ populations among total HIV RNA+ cells are indicated above each population. **B.** Infected cells with high and low levels of HIV transcripts can be identified regardless of implementation of HIV-seq. HIV RNA+ cells from PID8027 were colored according to whether they were analyzed using the conventional or HIV-seq pipelines for scRNA-seq analysis. **C.** The transcriptomes of $HIV_{low}$ and $HIV_{high}$ cells are similar. Shown is a UMAP depiction of $HIV_{high}$ cells represented as red dots and $HIV_{low}$ cells as pink dots, against a background of HIV RNA- CD4+ T cells depicted in gray. Shown are combined results for PID8026 and PID8027. **D.** $HIV_{high}$ and $HIV_{low}$ cells have similar transcriptomes, but $HIV_{high}$ cells exhibit decreased cell-surface CD4 protein expression. Shown are volcano plots displaying differentially expressed transcripts and proteins, for the two viremic donors harboring both $HIV_{high}$ and $HIV_{low}$ cells. Select up- and down-regulated transcripts/proteins are annotated. Red dots correspond to genes and proteins with $0.25\log_2$ fold-change expression and with adjusted p value $\leq 0.05$, as determined by Wilcoxon rank sum Test.

**Figure S4. Gating strategy for CD4+ T cell subset identification.** A sequential gating strategy was implemented using a combination of surface protein markers and transcripts. CD4+ T cells were defined as CD45+CD19-CD20-CD3+CD8- cells, to include HIV-infected CD4+ T cells that have downregulated cell-surface CD4. Classic CD4+ T cells subsets were then defined as follows: naïve (Tn: CD45RO-CD45RA+), central memory (Tcm: CD45RO+CD45RA-*CD27*+*CCR7*+), effector memory (Tem: CD45RO+CD45RA-CD27-CCR7-), transitional memory (Tm: CD45RO+CD45RA-*CD27*+*CCR7*-), memory CCR7+CD27- (CD45RO+CD45RA-*CCR7*+*CD27*-), regulatory T cells (Treg: CD45RO+CD45RA-*CD25*+IL7R-) and non-Tregs (CD45RO+CD45RA-*CD25*-IL7R- or *CD25*+IL7R+ or *CD25*-IL7R+) and T follicular helper (Tfh: CD45RO+CD45RA-*CXCR5*+PD1+) and non-Tfh (CD45RO+CD45RA-*CXCR5*-PD1- or *CXCR5*+PD1- or *CXCR5*-PD1+) cells. Shown are data from one representative participant (PID8027).

**Figure S5. All six clusters of memory CD4+ T cells from viremic PWH contain all classical memory CD4 subsets.** Shown are pie graphs depicting the proportion of classical memory CD4+ T subsets (Tcm, Tem, Ttm, and CCR7+CD27- memory cells) among the 6 clusters identified by scRNA-seq in specimens from viremic PWH.

**Figure S6. ART suppression elicits global changes in host transcriptomes.** Shown are UMAP plots depicting total CD4+ T cells from viremic (red) and suppressed (blue) time points for each of the three donors with paired specimens (PID1052, PID8026, and PID8027). For all three participants, CD4+ T cells during viremia localize in different regions of the UMAP relative to CD4+ T cells during ART suppression.

**Figure S7. Compared to CD4+ T cells during ART, CD4+ T cells during viremia present an activation of the integrated stress pathway with diminished ribosomal transcript expression levels.** Shown is a volcano plot displaying differentially expressed transcripts in CD4+ T cells from viremic vs. suppressed time points, with ribosomal transcripts and transcripts related to ISR pathway annotated. Red dots correspond to transcripts with 0.25$\log_2$ fold-change expression and with adjusted p value ≤ 0.05, as determined by the Wilcoxon rank sum test.
